# Sepsis Subtypes in Blood and Liver Define Precision Strategy for CXCR2 Blockade

**DOI:** 10.64898/2026.08.03.740897

**Authors:** Na Liu, Jenny Halbauer, Mohamed Albadry, Uta Dahmen, Nikolaus Gaßler, Brendon P. Scicluna, Michael Bauer, Adrian T. Press

## Abstract

Sepsis is classified into distinct transcriptomic endotypes. Yet specific cellular drivers for those subtypes remain ambiguous. Although dysregulated CXCL8–CXCR1/2 signaling and neutrophil hyperactivation are implicated in severe endotypes, the therapeutic window and organ-specific consequences of CXCR2 antagonism remain unclear. To determine whether murine transcriptomic subtypes (MTSs) recapitulate human consensus transcriptomic subtypes (CTSs) and evaluate how subtype state dictates the efficacy and trade-offs of Danirixin in polymicrobial sepsis. We reanalyzed septic patients whole-blood CITE-seq data to characterize CXCL8– CXCR2 signaling. Using a severity-stratified murine polymicrobial model, we evaluated Danirixin’s efficacy. Endpoints included 7-day survival, cytokine profiling, and hepatic histology. Early (24-h) multi-compartment assessments of bacterial burden, NETosis, immune infiltration, and paired blood–liver bulk RNA sequencing informed MTS classification and mechanistic insights. Human CTS1 exhibited neutrophil and progenitor expansion, elevated CXCL8/CD11b expression, and reduced CXCR2, defining a hyperactivated, dysregulated myeloid state. Applying CTS-derived gene signatures to murine tissues identified three MTSs correlating with pathogen load and interleukin-6. Human CTS1 dysregulation was mirrored in murine MTS3. Crucially, Danirixin improved 7-day survival in MTS3 sepsis. Conversely, in MTS1, Danirixin attenuated systemic NETosis and hepatic injury but exacerbated bacterial dissemination, without improving survival. Murine transcriptomic subtypes translationally model human sepsis subtypes. In severely dysregulated host-response states (MTS3/CTS1), CXCR2 antagonism effectively mitigates maladaptive, neutrophil-driven immunopathology. However, in less severe subtypes, it compromises early bacterial containment. These findings therefore support for endotype-guided precision targeting of CXCR2 in sepsis.

**One Sentence Summary:** Targeting CXCR2 improves sepsis survival only in a specific molecular subtype, showing that host traits dictate therapeutic outcomes.

## INTRODUCTION

Sepsis arises when a localized infection triggers a dysregulated systemic host response, multiorgan dysfunction, and death. Despite decades of investigation, broadly applied immunomodulatory therapies have failed, underscoring the biological heterogeneity of sepsis and the need for precision-based strategies (*1*). Recent large-scale transcriptomic studies have converged on three consensus transcriptomics subtypes (CTS1–3), which capture major axes of host-response endotpyes (*2*). CTS1 is characterized by hyperinflammation, including immature neutrophils and endothelial activation; CTS2 is associated with heme metabolism, coagulopathy and platelet-related programs; and CTS3 is defined by interferon signaling, lymphocyte and nonclassical monocyte features, and anticoagulant pathways. However, these endotypes are derived from high-dimensional blood transcriptomic data, and it remains unclear whether they can be translated into experimentally traceable, omics-free traits or approximated by routine clinical readouts.

Neutrophils are central to sepsis pathophysiology across host-response states and exemplify the dual nature of innate immunity in this syndrome. In hyperinflammatory settings, excessive neutrophil recruitment and neutrophil extracellular trap (NET) formation contribute to microvascular injury, thrombosis, and organ failure. Conversely, impaired neutrophil trafficking and antimicrobial function compromise early pathogen control and promote dissemination (*3–6*). Neutrophil mobilization and activation are predominantly regulated by the ELR^+^ chemokine– CXCR1/2 axis (*7*), while CXCR4 provides counter-regulating retention in the bone marrow (*8, 9*). In sepsis, sustained exposure to CXCR2 ligands drives receptor internalization and the emergence of CXCR2-low immature neutrophils, which are associated with sepsis severity, secondary infections and immunoparalysis (*10–13*).

Preclinical studies suggest that the therapeutic benefit of CXCR2 antagonism in sepsis is dependent on host-response context. In experimental septic peritonitis (*14, 15*), CXCR2 inhibition improves survival by limiting NETosis-driven lung injury, optimizing leukocyte recruitment kinetics, and promoting protective peritoneal CXCL10 expression. Beyond sepsis, CXCR2 antagonism has shown anti-inflammatory activity (*16, 17*). However, clinical experience in chronic obstructive pulmonary disease highlights the challenges of therapeutic targeting in biologically heterogeneous inflammatory diseases. Although CXCR2 antagonists reduce neutrophil activation, clinical responses are variable and overall benefit has been limited (*18, 19*).

Danirixin, a selective, reversible, competitive CXCR2 antagonist, is therefore an attractive translational candidate. We hypothesized that host-response heterogeneity, as captured by human CTSs and projected onto murine transcriptomic subtypes (MTSs), critically shapes the response to CXCR2 antagonism. We expected the therapeutic benefit to be confined to specific sepsis states where limiting pathological NETosis outweighs the cost of impaired early bacterial control. We further hypothesized that these effects could not be reflected in broad systemic leukocyte reprogramming, but rather in selective modulation of organ-specific injury, particularly in the liver. To test this, we combined a severity-stratified murine model of polymicrobial sepsis with longitudinal biomarker profiling, cross-organ assessment of bacterial burden, NETosis, immune infiltration, and transcriptomic analysis of paired blood and liver samples.

## RESULTS

### Single-cell profiling identifies myeloid dysfunction across consensus transcriptomic sepsis subtypes

We interrogated a public EMROx whole-blood CITE-seq dataset (*3*) from 26 septic patients stratified by CTS. PCA of global blood transcriptomic profiles demonstrated clear separation between healthy volunteers and CTS1–3 patients (**Fig. 1A**). Pseudobulk expression analysis identified CTS classifier genes (*2*) that distinguished sepsis subtypes from healthy volunteers (**Fig. 1B**), spanning glycolytic and metabolic regulators, erythroid and heme-associated genes, innate immune and inflammasome-related genes, and genes linked to cellular stress, RNA and protein regulation, and trafficking. A composite myeloid dysregulation score was elevated across all CTS groups (**Fig. 1C**). UMAP identified the major circulating immune cell populations (**Fig. 1D**); CTS1 showed a marked expansion of neutrophils and neutrophil progenitors, whereas CTS3 was enriched in lymphocyte populations (**Fig. 1E**).

**Fig. 1.**
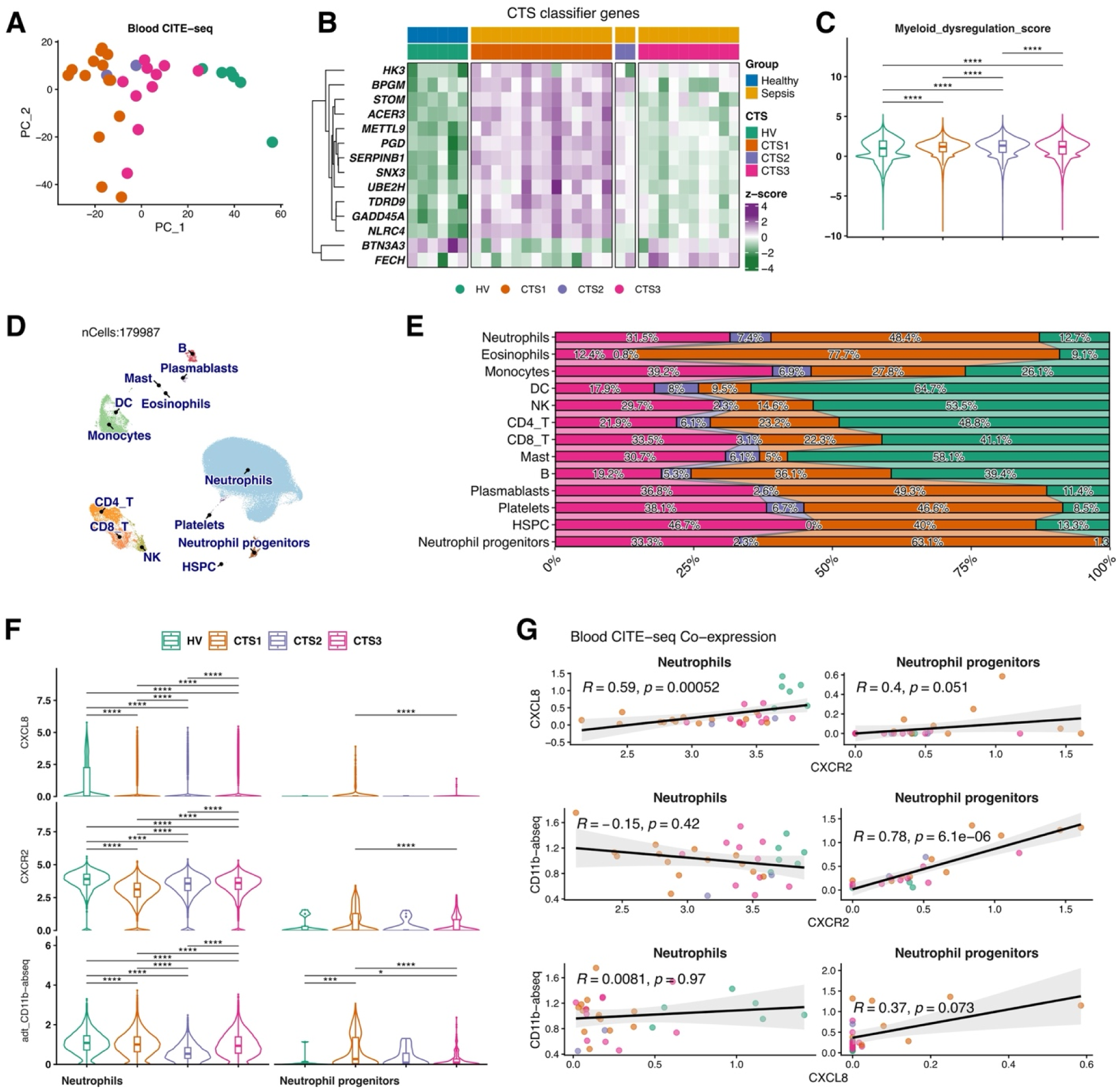
Single-cell profiling of circulating immune cells in human sepsis according to consensus transcriptomic subtypes (CTSs). (**A**) PCA of whole-blood CITE-seq profiles showing the separation of healthy volunteers (HV; n = 6) and sepsis patient subphenotypes CTS1 (n = 14), CTS2 (n = 2), and CTS3 (n = 10). (**B**) Heatmap showing the expression of CTS classifier genes across samples. (**C**) Myeloid dysregulation scores across all samples following log-transformation. (**D**) UMAP projection of annotated immune cell populations. (**E**) Relative proportions of cell clusters across CTSs. (**F**) Single-cell expression profiles of CXCL8 and CXCR2, together with surface CD11b protein levels, in neutrophils and neutrophil progenitors. (**G**) Spearman correlation analyses of CXCL8, CXCR2, and CD11b expression within the indicated neutrophils. Data are represented as violin plots showing the median ± IQR. Asterisks indicate statistical significance (ns, not significant; * p < 0.05, p < 0.01, *** p < 0.001, **** p < 0.0001).

Given the central role of the CXCL8–CXCR2 axis in neutrophils, we examined its expression across the neutrophil compartment. *CXCL8* transcripts and surface CD11b levels were elevated in neutrophils and progenitors across all CTS groups (**Fig. 1F**). In contrast, *CXCR2* was reduced in mature neutrophils most prominently in CTS1. The positive correlation between *CXCL8* and *CXCR2* transcript levels (R = 0.59, p = 0.0005;) in neutrophils and the correlation between surface CD11b abundance and *CXCR2* expression (R = 0.78, p < 0.0001) in progenitors confirmed this lineage-specific dysregulation in CTS1 (**Fig. 1G)**. These findings establish CTS1 as a neutrophil-dysregulation-dominant endotype characterized by hyperactivation, ligand excess, and receptor desensitization of the CXCL8–CXCR2 axis.

### Dose optimization establishes the pharmacodynamic basis for in vivo CXCR2 antagonism

Before evaluating Danirixin in sepsis, we defined its in vivo immunomodulatory profile in healthy mice (**Fig. S2A**). Dose escalation revealed a U-shaped response in blood neutrophil CXCR2 surface expression after hIL-8 stimulation (**Fig. S2B–E**), consistent with receptor saturation kinetics at higher doses. Based on these pharmacodynamic data, Danirixin at 15 mg kg ¹ was selected for all subsequent in vivo experiments.

### Blood and liver transcriptomics define compartment-specific host-response states with limited global perturbation by CXCR2 antagonism

To define the molecular basis of systemic and compartmentalized host responses, we performed bulk RNA sequencing of whole blood and liver from 38 mice 24 h after infection. Human CTS marker genes were translated to murine orthologs and applied as a classifier gene set to blood RNA-seq data from the PCI model, discriminating three murine transcriptomic subtypes (MTSs; **Fig. 2A**).

**Fig. 2.**
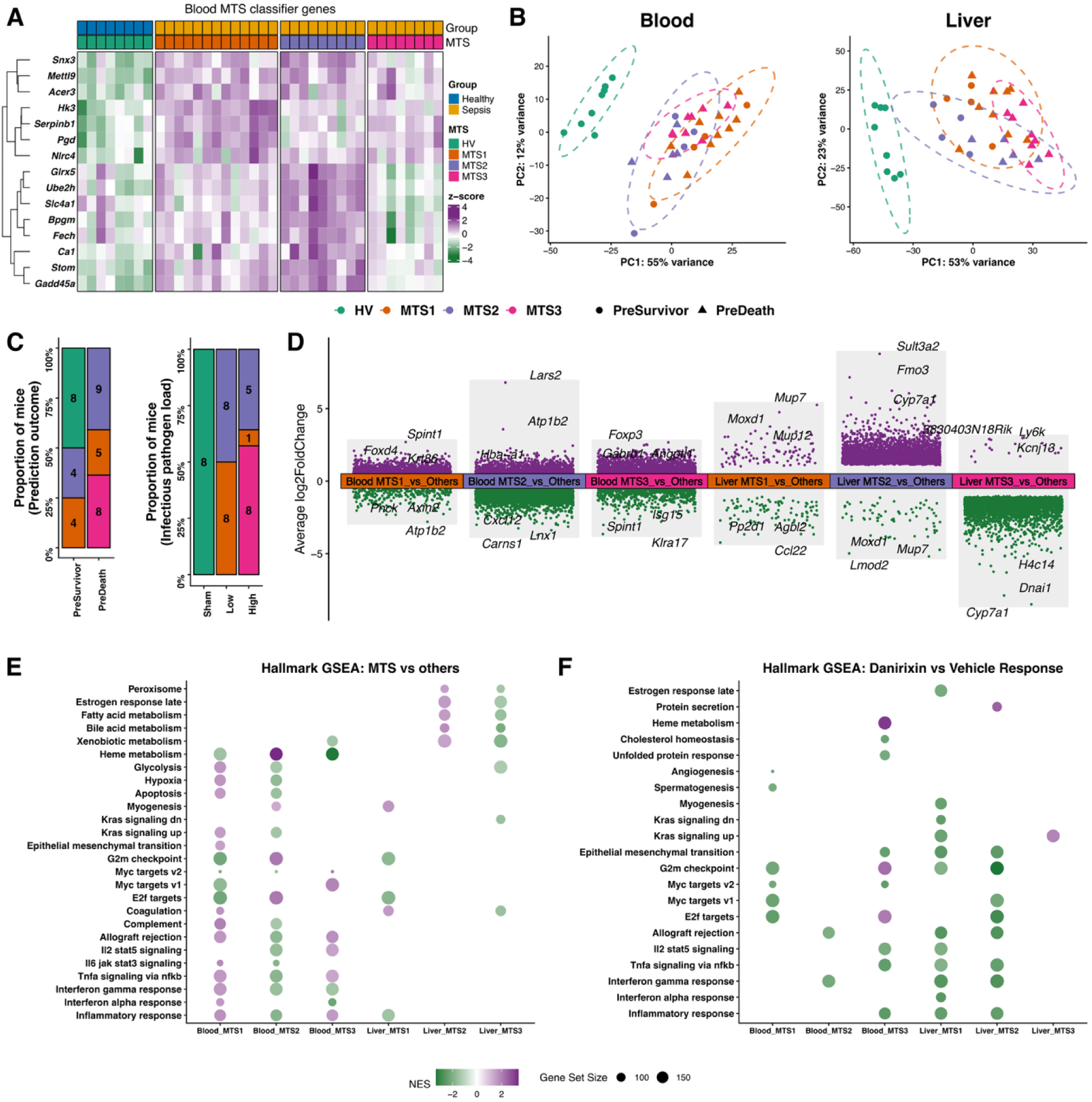
Murine Transcriptomic Subtyping (MTS) identifies distinct sepsis-associated states in the mouse blood and liver. (**A**) Heatmap illustrating the row-scaled expression of MTS classifier genes. The cohort comprises 38 blood samples in total: 8 HV, 14 MTS1, 8 MTS2, and 8 CTS3. (**B**) PCA plots demonstrating the distinct transcriptomic clustering in the blood and liver. Within the cohort, HV (4 Vehicle vs. 4 Danirixin) and MTS3 (4 Vehicle vs. 4 Danirixin) are balanced, whereas MTS1 (5 Vehicle vs. 9 Danirixin) and MTS2 (6 Vehicle vs. 2 Danirixin) show disparate distributions. (**C**) Distribution of MTSs faceted by predicted outcome and initial infectious pathogen load. (**D**) DGE among the MTS subgroups across the blood and liver. (**E**) Hallmark GSEA comparing MTS subgroups. (**F**) Hallmark GSEA illustrating the transcriptional response to Danirixin treatment versus vehicle.

In the polymicrobial sepsis model, survival differed by pathogen load, with rates of 93% in the low-load group and 70% in the high-load group (**Fig. S1A**). Plasma IL-6 levels at 24 h predicted 7-day outcome (AUC = 0.72; **Fig. S1B**). MTS classification, defined by combining pathogen load and 24-h IL-6, yielded AUC values of 0.86–0.95 across subtypes (**Fig. S1C**) and was applied to the full cohort. MTSs were major drivers of transcriptomic variance in both tissues (**Fig. 2B**). In blood, PC1 and PC2 explained 50% and 14% variance, separating MTSs, pathogen load, and predicted outcome. Danirixin treatment had no significant effect on either component. In liver, PC1 and PC2 accounted for 54% and 23% variance and stratified with MTSs, pathogen load, and outcome as dominant contributors. Treatment again had no effect on the liver transcriptomic structure.

Among the 38 paired blood and liver RNA-seq samples, MTSs differed markedly in transcriptional remodeling (**Fig. 2C–D**). In blood, MTS1 showed a stable transcriptome, with only 29 differentially expressed genes (DEGs) totally, whereas MTS2 and MTS3 exhibited extensive changes (MTS2: 1,847 downregulated and 949 upregulated DEGs; MTS3: 471 downregulated and 689 upregulated DEGs). In liver, MTS1 was transcriptionally quiescent (1 DEG), while MTS2 was dominated by induction (3,871 upregulated and 35 downregulated DEGs) and MTS3 by repression (997 downregulated and 0 upregulated DEGs). The opposing directionality of hepatic MTS2 and MTS3 responses highlights fundamentally distinct organ-level pathological states that are not captured by blood classification alone.

Gene set enrichment analysis (GSEA) placed these differences in a biological context (**Fig. 2E**). In blood, MTS1 was enriched for proinflammatory, interferon, and hypoxia programs. MTS2 was characterized by immune-metabolic reprogramming, and MTS3 by reduced interferon and increased cytokine signaling. In liver, MTS2 was distinguished by enrichment of fatty acid, bile acid, and xenobiotic metabolism, whereas MTS3 showed broad suppression of these hepatic programs.

Danirixin exerted modest transcriptomic effects (**Fig. 2F**). In blood, it selectively reduced inflammatory and proliferative signaling across subtypes. In liver, it attenuated interferon and stress-response programs in MTS1 and MTS2. Hepatic MTS classification, derived independently from liver transcriptomes, diverged from blood-based assignments (**Fig. S1D–G**), confirming that blood-derived endotype classifiers incompletely reflect organ-level pathology and that tissue context substantially modifies subtype expression.

### CXCR2 antagonism selectively alters compartment-specific neutrophil responses

Mice were assigned to MTSs based on inoculum burden and circulating IL-6 at 24 h before Danirixin evaluation in severity-stratified PCI sepsis. MTS3 represented the most severe phenotype (**Fig. 3A**), with higher clinical severity scores than MTS1 and MTS2 at both 6 h (both p < 0.0001) and 18 h after infection (vs. MTS1, p = 0.0002; vs. MTS2, p < 0.0001), lower body weight (vs. MTS1, p = 0.03; vs. MTS2, p = 0.01), and hypothermia at 18 h (vs. MTS1, p = 0.007; vs. MTS2, p = 0.02). Danirixin treatment did not alter these clinical parameters. Relative weights of the liver, lung, and spleen were also similar across MTSs, although MTS2 exhibited a higher spleen-to-body weight ratio than MTS1 (p = 0.009). Consistent with this severity gradient, MTS3 showed the highest IL-6 concentrations at 24 h across the peritoneum, blood, liver, and spleen (**Fig. 3B**). Danirixin had little effect on compartmental IL-6, except for a reduction in MTS1 spleen.

**Fig. 3.**
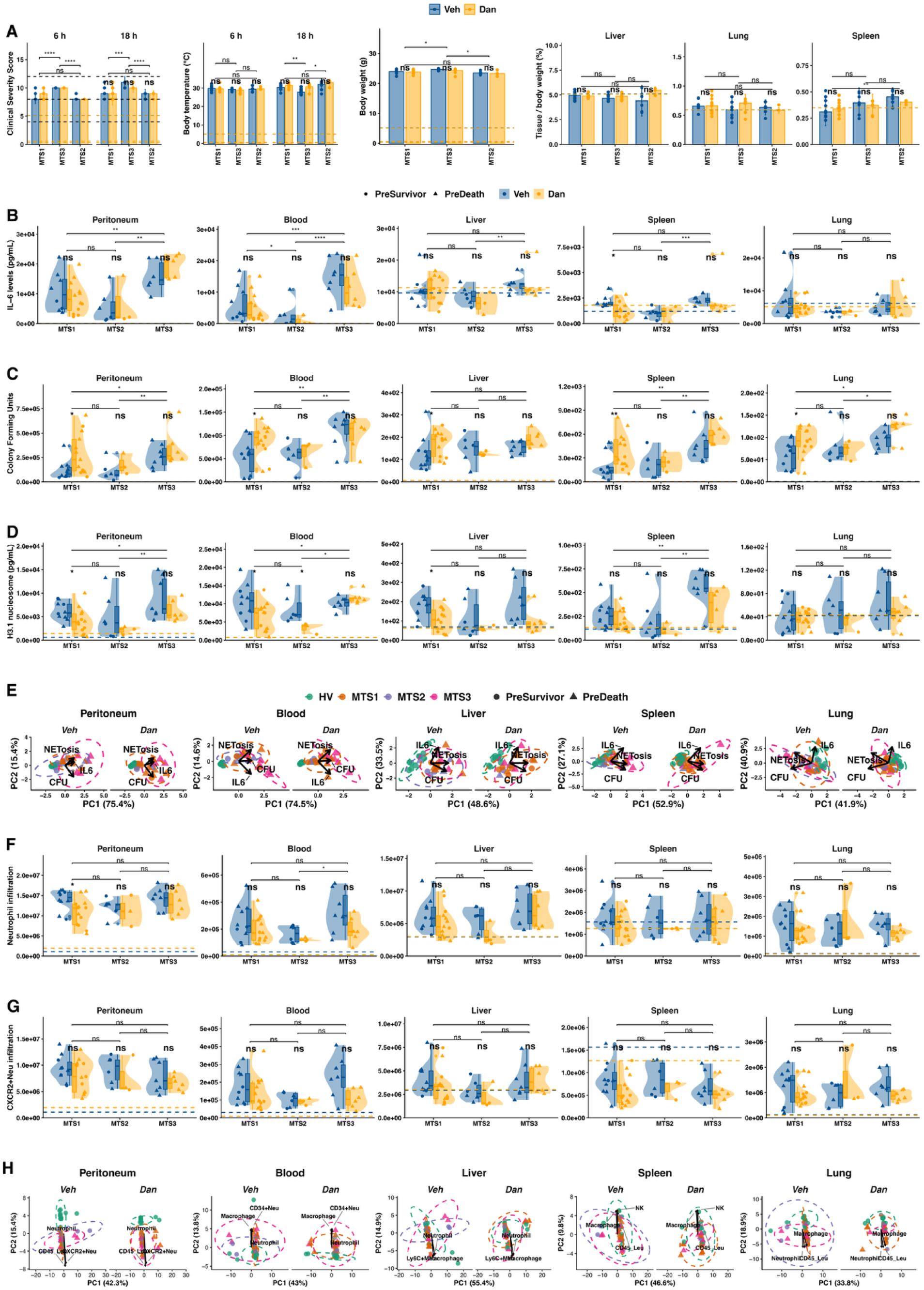
Multi compartment IL 6 concentrations, bacterial clearance, NETosis, and neutrophil dynamics at 24 h post infection. (**A**) Phenotypical characterization 24 h after infection. (**B-D**) Tukey boxplot for (B) IL 6 levels, (C) bacterial burden, and (D) histone 3.1 nucleosome concentration across compartments. Dashed lines depict sham levels. (**E**) PCA integrating data from B to D. (**F-G**) Flow cytometric quantification of (F) total neutrophil infiltration, and (G) CXCR2-expressing neutrophils within the corresponding tissues. (**H**) PCA of integrated cytometric cellular infiltrate data. Arrows indicate the top three cellular variables driving sample clustering. Overlaid boxplots indicate the median and interquartile range; whiskers extend to the minimum and maximum values. Asterisks denote statistical significance (ns: not significant; * p < 0.05, ** p < 0.01, *** p < 0.001, **** p < 0.0001).

Bacterial burden similarly tracked disease severity, with MTS3 exhibiting the highest pathogen counts in the peritoneum, blood, spleen, and lung, but not the liver (**Fig. 3C**). Critically, Danirixin increased bacterial burden in MTS1 across all five compartments: the peritoneum (p = 0.04), blood (p = 0.03), liver (p = 0.05), spleen (p = 0.01), and lung (p = 0.05), with no detectable effect in MTS2 or MTS3, indicating that CXCR2 antagonism compromises early pathogen containment specifically in less severe disease states. NETosis also followed a compartment- and severity-dependent pattern. MTS3 showed the highest NETosis in the peritoneum, blood, and spleen (**Fig. 3D**), whereas NETosis in the liver and lung were comparable across MTSs. Danirixin reduced NETosis of MTS1 in the peritoneum (p = 0.03), blood (p = 0.05), and liver (p = 0.02), and of MTS2 in the blood (p = 0.02). The combination of increased bacterial burden and reduced NETosis in MTS1 following Danirixin treatment indicates disruption of NET-mediated bacterial containment in this subtype.

Biomarker correlations across compartments confirmed that IL-6, CFU, and NETosis were tightly coupled in the peritoneum and blood (R = 0.68–0.78, all p < 0.0001), with moderate associations in the lung, liver, and spleen (**Fig. S3A–B**). Integrated PCA of IL-6, bacterial burden, and NETosis confirmed that pathogen load and predicted outcome, but not Danirixin, structured the multi-organ pathological landscape across all compartments (all p < 0.001; **Fig. 3E**). Flow cytometry revealed that neutrophil accumulation and CD45 infiltration tracked disease severity across organs (**Fig. S4A**).

Total neutrophil accumulation in the blood was higher in MTS3 than in MTS2 (p = 0.04), and CD45 immune cell levels were highest in the blood of MTS3 (**Fig 3F)**. Danirixin did not alter total neutrophil infiltration across compartments, except for a reduction in the peritoneum of MTS1 (p = 0.03). Consistent with functional rather than numerical modulation, Danirixin did not reduce the accumulation of CXCR2 neutrophils across organs within any MTS group (**Fig. 3G**), but it decreased neutrophil CD11b expression in the peritoneum of MTS3 (p < 0.05; **Fig. S4C**), indicating selective attenuation of neutrophil activation state rather than recruitment per se. In MTS2, Danirixin reduced the elevated CD45 immune cell levels in the blood (**Fig. S4B**), and in MTS1, macrophage abundance was reduced in the spleen (**Fig. S4D**), suggesting a shift away from innate myeloid dominance in less severe disease states.

Global PCA of tissue immune profiles supported these localized effects. MTSs separated samples in the peritoneum, blood, and lung (all p < 0.001 to 0.05; **Fig. 3H**), while Danirixin had minimal effects on the global immune landscape, except in the spleen, where treatment shifted samples along PC2 (19% variance explained, p = 0.02), driven primarily by CD45^+^ leukocytes, macrophages, and NK cells.

### CXCR2 antagonism confers a sepsis subtype effect on mortality or liver injury

Over 7 days, Danirixin did not alter clinical severity scores, body weight, temperature, or systemic CXCL1/CXCL2 levels (**Fig. 4A–C**). It exerted subtype- and time-dependent effects on circulating IL-6, increasing levels in MTS1 at 24 h (p = 0.006) and reducing them in MTS2 at 48 h (p = 0.02; **Fig. 4D**). Despite comparable clinical trajectories, Danirixin improved survival in MTS3, without notable effects in MTS1 or MTS2 (**Fig. 4E**). Accordingly, survival was not altered in MTS1 or MTS2.

**Fig. 4.**
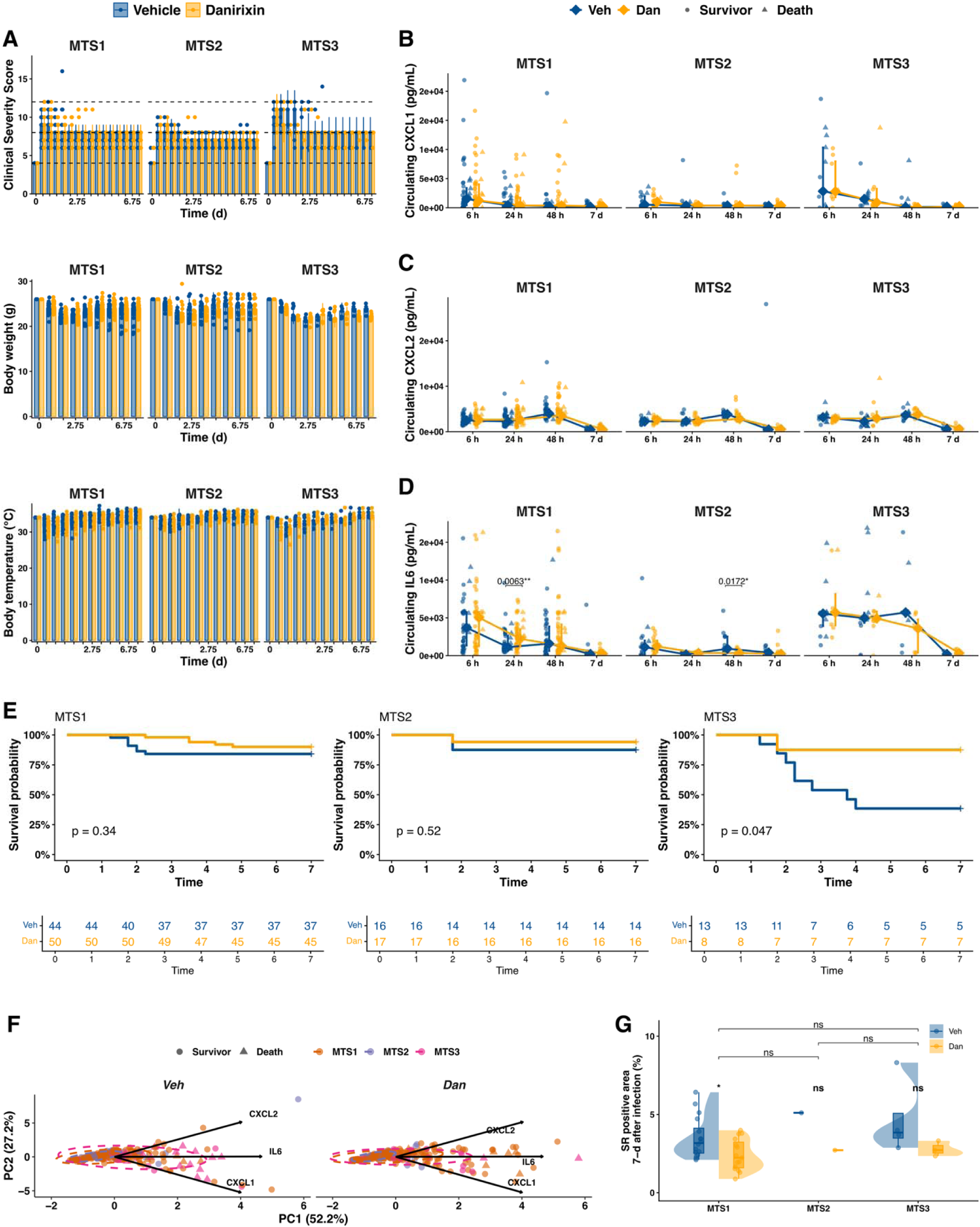
Clinical severity, survival, circulating inflammatory mediators, and hepatic collagen deposition in a sepsis model following Danirixin treatment. (**A**) Longitudinal clinical severity score, body weight, and temperature over 7 d after infection. (**B–D**) Time resolved circulating concentrations of CXCL1/2 and IL 6 over the 7 d observation. (**E**) Kaplan–Meier survival curves and numbers at risk over 7 d, stratified by predicted MTS group from a retrospective survival analysis. (**F**) PCA integrating longitudinal CXCL1/2 and IL 6 measurements. (**G**) Hepatic collagen deposition assessed 7-d after infection, quantified as histologically identified Sirius Red positive area. Individual data points represent single animals; Tukey boxplot with a bar at the median and whiskers spanning the interquartile range. Asterisks indicate statistical significance (ns: not significant, * p < 0.05).

Biomarker predictive performance differed according to pathogen load (**Fig. S5A**). CXCL1 was the strongest predictor of outcome in the low-load group (AUC up to 0.93 at 48 h), while IL-6 outperformed chemokines in the high-load group (AUC up to 0.83). In MTS1, CXCL1 distinguished non-survivors from survivors across all time points (6 h, 24 h, 48 h; all p ≤ 0.04); in MTS3, IL-6 was the dominant prognostic marker (p = 0.008 at 24 h; **Fig. S5B–C**).

PCA of serial blood cytokine measurements collected from 6 h to 7 days after infection confirmed IL-6 as the dominant contributor to inflammatory trajectory (PC1, 52% variance), associated with MTS, pathogen load, and 7-day survival (all p < 0.0001), but independent of Danirixin treatment (**Fig. 4F**). Sepsis-induced liver injury was quantified by collagen-positive area on whole-slide images (**Fig. S6A**).

IL-6 correlated with hepatic collagen deposition across time points (R = 0.5, p = 0.001; **Fig. S6B**), linking systemic inflammation to organ fibrosis. Danirixin reduced hepatic collagen in MTS1 (p = 0.014; **Fig. 4G**) but not in MTS2 or MTS3, suggesting that CXCR2 antagonism selectively modulates long-term liver remodeling in less severe subtypes. This notion is consistent with the anti-NETotic and anti-inflammatory effects observed at 24 h in this subtype yet uncoupled from any survival benefit.

## DISCUSSION

Human single-cell data identified neutrophil activation and dysregulated CXCL8–CXCR2 signaling as central features of the sepsis response. Across human consensus transcriptomic subtypes (CTSs), sepsis was broadly characterized by neutrophil dysregulation. CTS1 exhibited an expansion of neutrophils and their progenitors, along with elevated *CXCL8* and CD11b, and reduced *CXCR2* in mature neutrophils. This supports the concept that sustained inflammation drives neutrophil hyperactivation and receptor desensitization, positioning the CXCL8–CXCR2 axis at the core of severe, myeloid-dominant sepsis (*3, 20*). Critically, this neutrophil-dysregulation signature in human CTS1 was mirrored in MTS3, establishing a cross-species biological parallel that anchors the translational relevance of our experimental model.

The murine host response recapitulated major axes of human CTS biology, supporting the view that MTSs represent a translational extension of human CTSs (*2, 21*) rather than independent entities. The neutrophil-predominant, dysregulated human CTS1 closely resembled MTS3, which was characterized by high pathogen load, elevated IL-6, and extensive transcriptional remodeling in the blood and liver. Conversely, human CTS3 aligned with the less severe MTS1. Rather than claiming direct functional equivalence, this alignment demonstrates that human and murine transcriptomic frameworks capture analogous positions along a shared disease continuum, from mild immune perturbation in MTS1/CTS3 to severe innate-driven dysregulation in MTS3/CTS1. These findings suggest that disease severity and inflammatory trajectory, rather than species differences, are the primary determinants of subtype identity.

Paired blood and liver transcriptomics revealed that subtype biology is compartmentalized. While blood MTS assignment captured major systemic variation, hepatic transcriptomes emphasized metabolic adaptation, repression of hepatic programs, tissue injury, and direct CTS classification of liver RNA-seq diverged from blood-based classification. This divergence reflects the liver operating as a semi-autonomous immunometabolic compartment whose response to both sepsis and CXCR2 antagonism is shaped by local signals (*22*). Accordingly, Danirixin induced minimal transcriptomic changes in blood yet profoundly altered hepatic pathways, attenuating interferon programs and preserving metabolic pathways such as fatty acid and bile acid metabolism, which were otherwise suppressed in untreated sepsis. These findings demonstrate that the therapeutic efficacy of immunomodulatory agents cannot be inferred from systemic biomarkers alone.

These subtype- and compartment-specific differences fundamentally shape the therapeutic window and effects of CXCR2 antagonism. In MTS1, where pathogen burden remains containable, CXCR2-driven neutrophil recruitment appeared to be the critical determinant of outcome. Danirixin impaired this response, increasing bacterial dissemination across all five compartments without conferring survival benefit, and its anti-NETotic and hepatoprotective effects at 24 h failed to translate into improved 7-day outcome. These findings demonstrate that organ-level protection and survival can be dissociated when bacterial containment is compromised. In MTS2, the intermediate inflammatory and infectious burden placed the host in a therapeutic null zone, in which Danirixin attenuated NETosis and cytokine levels without shaping either the bacterial-clearance or immunopathology axis sufficiently to alter survival. In MTS3, overwhelming pathogen dissemination rendered neutrophil-mediated bacterial clearance insufficient regardless of treatment, shifting the dominant determinant of mortality to lethal systemic NETosis, the precise target of CXCR2 antagonism. Accordingly, Danirixin attenuated this response and improved 7-day survival without further compromising bacterial control.

Danirixin’s selective, reversible, and competitive pharmacological profile appears central to this subtype-dependent efficacy. It reduced systemic NET release without broadly suppressing the cytokine response, uncoupling neutrophil-driven tissue injury from systemic inflammation (*14, 23, 24*). Competitive antagonism limits excessive ligand-driven neutrophil activation while avoiding the complete functional paralysis observed in genetic knockout models, explaining why Danirixin preserved sufficient antimicrobial function in MTS3, while dampening protective NET-mediated containment in MTS1 (*25, 26*). Although receptor trafficking was not directly measured, the selective reduction in NETosis without global neutrophil suppression is consistent with the CXCR2 downregulation observed in septic patients (*13, 27*).

Temporal immune profiling confirmed that Danirixin acts as a selective immunomodulator rather than a broad immunosuppressant. It reduced CD11b surface expression in MTS3 peritoneum, without reducing CXCR2 neutrophil accumulation, indicating attenuation of activation state rather than recruitment per se (*15*). Reduced splenic macrophage abundance in MTS1 further suggested a shift away from pathogenic granulocytic dominance, reminiscent of myeloid reprogramming observed following CXCR2 inhibition in tumor models (*28–30*). Moreover, Danirixin reduced hepatic collagen deposition in MTS1 at 7 days, indicating that CXCR2 antagonism influences long-term liver remodeling, although this organ-protective effect remained dissociated from any survival benefit in this subtype (*31, 32*). Together, these findings position the liver as a primary site of therapeutic modulation by CXCR2 antagonism, potentially involving both canonical and non-canonical signaling pathways beyond neutrophil chemotaxis (*33–37*).

Several limitations warrant consideration. Our polymicrobial model captures core sepsis features but may not fully recapitulate human heterogeneity, and varying pathogen loads may confound severity-group comparison. The MTS3 survival finding was derived from a post-hoc subgroup of n = 8 Danirixin-treated animals. Although this reflects the natural severity distribution after transcriptomic classification within a well-powered overall survival cohort (n = 55) and is mechanistically supported by converging 24-h endpoint data, independent replication in severity-enriched cohorts is required.

MTS assignment inherently integrated baseline host response. Human CTSs and MTSs, though biologically correspondent, are not functionally interchangeable across species. Fixed dosing intervals may not align with the optimal therapeutic window, and 24-h and 7-day endpoints limit insight into hyper-acute dynamics and longer-term recovery. Prospective subtype-guided stratification and dynamic treatment windows are needed to define the clinical conditions under which CXCR2 antagonism can be safely deployed.

This study positions the CXCL8–CXCR2 axis as a central, subtype-dependent determinant of sepsis outcomes across humans and mice. Human CTSs were compared to a murine framework, demonstrating that host-response heterogeneity and tissue context govern the effects of CXCR2 antagonism. Application of a CTS-derived gene panel to murine transcriptomes identified three MTSs that tracked with pathogen load and plasma IL-6 levels. Danirixin improved 7-day survival in MTS3, whereas in MTS1 it reduced systemic NETosis and hepatic injury but increased bacterial dissemination, without conferring a survival benefit.

## MATERIALS AND METHODS

### Re-analysis of consensus transcriptomic subtype classification in human single-cell data

Publicly available whole blood cellular indexing of transcriptomes and epitopes by sequencing (CITE-seq) (*3*) from 26 septic patients and 6 healthy volunteers (HV) were obtained and re-analyzed. Single-cell processing, principal component analysis (PCA), cell-type annotation, and visualization were conducted using Seurat (R v5.0.1) (*38*) and SeuratExtend (R v1.2.0) (*39*).

Pseudobulk RNA-seq profiles were generated by aggregating raw gene counts across all cells from each individual using the AggregateExpression function. Pseudobulk count matrices were normalized using the trimmed mean of M-values method in edgeR (R v4.8.2). Lowly expressed genes were removed using filterByExpr, and gene expression values were transformed to log2 counts per million (CPM) using cpm(log = T, prior.count = 1). For unsupervised analyses, top 80% most variable genes were selected on the basis of median absolute deviation. Gene symbols were mapped to Ensembl gene identifiers using org.Hs.eg.db (R v3.22.0).

Consensus transcriptomic subtypes (CTSs) were assigned using the run_subtype_classifier function from ctsSubtypeR (R v0.0.0.9000). Normalized pseudobulk expression profiles from septic patients were compared with those from HVs using a random forest classifier trained on the reference sepsis dataset with 500 trees. Each sepsis sample was assigned to one of three CTS subtypes.

To quantify myeloid dysfunction, Seurat log-normalized expression values were first linearized by exponentiation. A dysfunction score was then calculated as the geometric mean of a curated detrimental gene set minus the geometric mean of a protective gene set, as described by Moore et al (*21*). Expression dynamics of key immune mediators were assessed using transcript abundance of *CXCR2* and *CXCL8*, together with surface protein expression of CD11b measured by AbSeq. To evaluate multimarker co-expression patterns, patient-level pseudobulk averages were calculated for selected myeloid cell populations and analyzed using Spearman’s rank correlation.

### Animal experiments

All animal experiments were conducted in accordance with German legislation and were approved by the local animal welfare authority (Thuringian State Administrative Office, Thuringia, Germany; approval no. UKJ-23-003). Use of human fecal slurry prepared from healthy volunteers for induction of sepsis was authorized by the Ethical Committee of Friedrich-Schiller University Jena (approval no. 2019-1413-Material). The microbial composition of the infectious inoculum has been described (*40, 41*).

Sex-balanced C57BL/6J mice aged 12–16 weeks were used throughout. After a 2-week acclimatization, animals were housed under specific pathogen-free conditions in individually ventilated cages (5 mice per cage) at 22°C, 45% ± 5% relative humidity, and a 12 h light/dark cycle, with food and water were available ad libitum. The study design followed the Minimum Quality Threshold in Preclinical Sepsis Study (MQTiPSS) recommendations.(*42*) Animals were randomized to experimental groups on a cage basis. Investigators performing sepsis induction were blinded to subsequent treatment. Reporting followed the *Animal Research: Reporting In Vivo Experiment* (ARRIVE) guidelines, which were provided in the supplement (*43*).

Three complementary animal experiments were performed to define pharmacodynamic effects, immunological consequences, and therapeutic efficacy of CXCR2 inhibition. First, a dose-response experiment was conducted in healthy mice to characterize baseline pharmacodynamic effects of Danirixin. Second, an acute endpoint study was performed 24 h after induction of polymicrobial sepsis to assess early immunological and microbiological effects of treatment. Third, a 7-d survival study was conducted to determine the effect of CXCR2 inhibition.

To increase clinical relevance, the model incorporated human-relevant polymicrobial pathogens for abdominal sepsis, analgesic treatment with Metamizole, first-line antibiotic therapy with Meropenem, and fluid resuscitation with Ringer solution. Disease severity was monitored using a validated clinical scoring system and IL-6-based survival prediction (*44–46*).

### Dose-dependent CXCR2 inhibition in healthy mice

A total of 60 healthy mice received intraperitoneal (i.p.) Danirixin at doses of 5, 15, or 30 mg kg^-1^. Danirixin (GSK1325756; MedChemExpress, #HY-19768, USA) was prepared in DMSO as 30 mg mL^-1^ stock solution and diluted freshly in phosphate-buffer solution (PBS) immediately before injection. Vehicle-treated and untreated naïve mice were served as controls.

At 1 h after injection, peritoneal lavage fluid (PLF) and cardiac whole blood were collected and incubated for 1 h at 37°C with human IL-8 (10 nmol L^-1^) or PBS. Samples were then placed on ice, centrifuged at 600 g for 5 min at 4°C, and resuspended in staining buffer (BioLegend, #420201, USA). After Fc receptor blockade (Miltenyi Biotec, #130-092-575, Germany) for 10 min, cells were stained for 30 min at 4°C in the dark using fluorophore-conjugated antibody cocktail containing PE anti-CD45 (BioLegend, #103106, USA), BV421 anti-F4/80 (BioLegend, #123132, USA), APC anti-Ly6G (BioLegend, #127614, USA), PC5.5 anti-CD11b (BioLegend, #101228, USA), and FITC anti-CXCR2 (BioLegend, #149310, USA). Cells were washed, resuspended in the staining buffer and analyzed immediately on a CytoFLEX flow cytometer (Beckman Coulter, USA).

Flow cytometry data were analyzed using FlowJo v10 (BD Bioscience). Surface expression of CXCR2 and CD11b was quantified as geometric mean fluorescence intensity (gMFI) in defined myeloid subsets. Cell subsets were identified as neutrophils (CD45 CD11b Ly6G), monocytes (CD45 CD11b Ly6C), and macrophages (CD45 CD11b F4/80 ; **Fig. S2A**).

### Acute 24 h study of CXCR2 inhibition in polymicrobial sepsis

The early immunological effects of Danirixin were assessed 24 h after infection. A total of 72 mice were allocated by cage to six experimental groups (n = 12 per group) across six independent repetitions: (i) no pathogen load + vehicle, (ii) no pathogen load + Danirixin, (iii) low pathogen load + vehicle, (iv) low pathogen load + Danirixin, (v) high pathogen load + vehicle, and (vi) high pathogen load + Danirixin.

Polymicrobial sepsis was induced using the established peritoneal contamination and infection (PCI) model (*40, 41*). Mice received an i.p. injection of either 0 or 60 (1.2 × 10^11^ bacterial counts; low-pathogen load sepsis) or 100 µL (2 × 10^11^ bacterial counts; high-pathogen load) of standardized human fecal slurry in Ringer solution. All mice received supportive therapy with oral Metamizole (2.5 mg per mouse, four times daily) and subcutaneous Meropenem (25 mg kg^-1^ body weight, twice daily). Clinical severity scores (CSS) were recorded four times daily, body weight was measured once daily, and perianal surface temperature was assessed twice daily using an infrared thermometer (FLUKE, USA) (*47*).

Danirixin (15 mg kg^-1^) or vehicle (20% DMSO in PBS) was administered as a single i.p. injection 6 h after infection. At 24 h, all mice were anesthetized with 2.5% isoflurane and sampled. PLF was collected first by aseptically exposing the abdominal cavity, instilling 2 mL sterile PBS containing 5 mmol L^-1^ EDTA, gently massaging the abdomen, and aspirating the fluid. Cardiac whole blood was then collected, followed by excision of the lung, liver, and spleen tissues for downstream analyses.

### Seven-day survival study of CXCR2 inhibition in polymicrobial sepsis

Therapeutic efficacy was assessed in a 7-d survival study comprising 148 mice subjected to PCI-induced polymicrobial sepsis. Sepsis induction and supportive treatment were performed as described for the 24 h experiment and continued until the 7-d endpoint. Danirixin was administered i.p. at 15 mg kg^-1^ at 6, 30, and 54 h after infection. Vehicle-treated mice received an equivalent volume. Peripheral blood was collected longitudinal from the tail vein at 6, 24, and 48 h after infection into 180 µL EDTA-PBS solution (10 nmol L^-1^) (*48, 49*).

At day 7, surviving mice were euthanized by i.p. overdosed of ketamine/xylazine, followed by terminal blood collection via cardiac puncture. Whole blood was centrifuged at 1,000 g for 10 min to isolate plasma, which was aliquoted and stored at -80°C for subsequent cytokine and immunological analyses (*44, 50*). Liver and spleen were harvested, and tissue samples were processed into single-cell suspensions for flow cytometry.

### Tissue preparation, RNA sequencing, and analysis

Whole blood and liver tissues were collected from 38 mice at 24 h after infection, snap-frozen in liquid nitrogen, and stored at -80°C until processing. Total RNA from whole blood was extracted using Quick-RNA™ Whole Blood kit (Zymo Research, #R1201, USA) according to the manufacturer’s instructions. Ribosomal and globin RNA depletion was performed by Novogene GmbH (Munich, Germany). Total RNA from the liver tissues was isolated by Novogene using standard phenol-chloroform extraction. RNA quality and integrity were assessed using an Agilent 5400 Bioanalyzer and samples with an RNA integrity number ≥ 6 were used for library preparation. Poly(A)-enriched mRNA libraries were prepared by Novogene, and paired-end sequencing (2 × 150 bp) was performed on an Illumina NovaSeq 6000 platform across multiple lanes.

Raw FASTQ files were quality filtered and adapter trimmed using Trimmomatic (v0.40) (*51*). Filtered reads were aligned to the Mus musculus reference genome (GRCm39) using STAR (v2.7.11b) (*52*). Gene-level counts were generated with featureCounts from Subread (v2.0.6) (*53*) using Ensembl mouse gene annotation (v115) (*54*). Genes detected in fewer than 75% samples were excluded. Batch effects related to experimental run were corrected using ComBat-seq from sva (R v3.58.0) (*55*). Normalization, variance-stabilizing transformation, and differential gene expression analysis were conducted using DESeq2 (R v1.48.2) (*56*).

### Murine transcriptomic subtype classification in murine bulk RNA-seq data

Blood RNA-seq data from 38 mice were used to assign CTS previously defined in human sepsis (*2*). Mouse gene symbols were converted to human orthologs using orthogene (R v1.19.3) (*57*) followed by mapping to Ensembl gene identifiers using org.Hs.eg.db (R v3.22.0). Murine transcriptomic subtype (MTS) classification was performed using ctsSubtypeR (*2*). PCA was performed separately for blood and liver using variance-stabilized counts generated with DESeq2 (R v1.50.2) (*56*). Classifier gene expression was displayed as z score-scaled heatmaps using ComplexHeatmap (R v2.26.1) (*58*).

To identify subtype-specific transcriptional programs, one versus rest differential expression analyses were conducted for each MTS by comparing one subtype from the other two subtypes. Comparisons were performed for the blood and liver using raw count matrices in DESeq2. Genes with fewer than one count across 80% samples were excluded. Genes with a BH adjusted p value < 0.05 and an absolute log fold change (log FC) > 0.5 were considered differentially expressed genes (DEGs) and visuliazed with scRNAtoolVis (R v0.1.0).

Pre-ranked gene set enrichment analysis (GSEA) was performed using fgsea (R v1.36.2) (*59*) against the MSigDB (*60*) Hallmark gene set collection. For both MTS-versus-others and treatment-within-MTS comparisons, genes were ranked using the metric −log_10_(p value) × sign(log FC). Ranked gene lists were analyzed separately for each tissue and comparison. Pathways with a BH-adjusted p-value < 0.05 were considered significantly enriched and visualized using clusterProfiler (R v4.18.4) (*61*).

### Preparation of single-cell suspensions

PLF was passed through a 70 µm cell strainer (Corning, #431751, USA) and centrifuged at 500 g for 5 min at 4°C. The resulting cell pellet was washed once and resuspended in the staining buffer. To isolate leukocytes from whole blood, 50 µL freshly collected blood was mixed with 150 µL PBS and 200 µL 3% dextran solution. The mixture was incubated at room temperature for 20 min to allow red blood cell (RBC) sedimentation. The leukocyte-rich supernatant was then collected, centrifuged at 500 g for 5 min, washed once, and resuspended in the staining buffer.

Liver, lung, and spleen tissues were harvested, mechanically dissociated, and passed through a 70 µm nylon cell strainer to obtain single-cell suspensions. Liver homogenates were subjected to an initial low-speed centrifugation step at 50 g to pellet hepatocytes. The resulting non-parenchymal cell (NPC)-enriched supernatant was collected and centrifuged again at 500 g for 5 min. Lung and spleen homogenates were directly centrifuged at 500 g for 5 min. Cell pellets from all tissues were treated with RBC lysis buffer (BioLegend, # 420302, USA), washed, and finally resuspended in the staining buffer at a concentration of 10^6^ cells per 100 μL for subsequent flow cytometric analysis.

### Flow cytometry

Cell pellets isolated from the different organs were resuspended in Fc receptor blocking solution and incubated for 10 min at 4°C. Cells were then stained for 30 min at 4°C in the dark with fluorochrome-conjugated antibody cocktails. The antibody panel comprised BV421 anti-CD146 (Miltenyi Biotec, #130-103-378, 1:50, Germany), BV605 anti-Ly6G (BioLegend, #127639, USA), BV650 anti-F4/80 (BioLegend, #123149, USA), BV785 anti-CD3 (BioLegend, #100232, USA), FITC anti-citH3 (BD Biosciences, #558610, USA), PerCP/Cy5.5 anti-CD45 (BioLegend, #103132, USA), PE anti-CXCR2 (BioLegend, #149304, USA), PE-CFS594 anti-CD34 (BioLegend, #128616, USA), PE-Cy5 anti-CD11b (BioLegend, #101210, USA), PE-Cy7 anti-CD49b (BioLegend, #108992, USA), APC anti-MPO (BD Biosciences, #570233, USA), AF700 anti-Ly6C (BioLegend, #128024, USA), APC-Cy7 anti-CD19 (BioLegend, #152412, USA). After staining, cells were washed once and resuspended in 200 µL staining buffer for acquisition. Flow cytometry data were acquired immediately on a FACSymphony A1 (BD Biosciences, USA) and analyzed using FlowJo.

Fluorescence compensation and instrument voltage settings were optimized according to standard laboratory procedures (*62*). Cell populations were identified from exported raw .fcs files using the following gating strategy. Viable singlets were selected based on forward and side scatter (FSC/SSC) characteristics. Myeloid cells were defined as CD45 CD11b ; neutrophils as CD45 CD11b Ly6G Ly6C ^/^ ; macrophages as CD45 CD11b^+^ F4/80 ^/^ Ly6C Ly6G ; monocytes as CD45 CD11b Ly6G Ly6C ; T cells as CD45 CD11b CD19 CD3 ; B cells as CD45 CD11b CD19 CD3 ; and natural killer (NK) cells as CD45 CD11b ^/^ CD49b . Neutrophil extracellular trap (NET) formation was quantified as MPO citH3 events within the neutrophil gate. The relative frequency (%) of each population and gMFI values were compared between experimental groups (**Fig. S5A**).

### Enzyme-linked immunosorbent assay (ELISA)

Circulating cytokine levels at 6, 24, 48 h, as well as 7 d after infection, were quantified using a modified sequential ELISA (*44, 63*). Plasma concentration of IL-6 (BioLegend, ELISA MAX Set Mouse IL-6, #431301, USA), CXCL1 (Invitrogen, #900-K127, USA), and CXCL2 (Invitrogen, #900-K152, USA) were measured according to the manufacturers’ instructions. Optical densities were recorded using an EnSpire multimode plate reader (PerkinElmer, USA). For IL-6, absorbance was measured at 450 nm with background correction at 570 nm. For CXCL1/2, absorbance was measured at 405 nm with correction at 650 nm. All samples were analyzed in duplicate, and cytokine concentrations were calculated using a four-parameter logistic standard curve.

Circulating nucleosomes containing histone H3.1 were quantified using the Nu.Q® Discovery H3.1 ELISA kit (CE-IVDD, Belgian Volition SRL, Isnes, Belgium) according to the manufacturer’s protocol. Measurements were performed on plasma and tissue homogenates collected at 24 h after infection. Optical density was measured at 450 nm using the EnSpire multimode plate reader.

### Bacterial quantification

Bacterial burden in collected samples was determined using colony-forming unit (CFU) assay. Cell-free suspensions were prepared from PLF, whole blood, lung, liver, and spleen samples. For each sample, 30 µL suspension was diluted 1:10 in 270 µL sterile DPBS. For each dilution, 100 µL was plated in triplicate on plate count agar (Carl Roth, #X930.2, Germany). Plates were incubated at 37°C for 48 h, and visible colonies were then counted. Bacterial load was calculated and expressed as CFU per microliter (CFU µL^-1^) for fluid samples and CFU per gram of tissue (CFU g^-1^) for solid organs.

### Tissue processing and histological analyses

Liver tissues collected at 24 h and 7 d were fixed in 4% paraformaldehyde (PFA) at 4°C for 24 h. Fixed samples were subsequently processed for formalin-fixed, paraffin-embedded (FFPE) preparation using a Leica TP1020 tissue processor (Leica, Germany). The processing protocol comprised sequential dehydration, clearing, and paraffin infiltration steps as follows: 40% ethanol for 30 min; 70% ethanol for 30 min; 96% ethanol for 45 min (twice); 100% ethanol for 45 min followed by 40 min; xylene for 60 min (twice); and paraffin infiltration for 120 min (twice). Tissues were then embedded in paraffin blocks and sectioned at 3 µm using a Leica RM2165 microtome (Leica Biosystems, Germany).

FFPE liver sections collected at 24 h and 7 d after infection were subjected to hematoxylin and eosin (H&E) staining using a standard protocol in the pathology routine laboratory at Jena University Hospital. Briefly, sections were fixed in formalin for 10 min, dehydrated in isopropanol, and stained with Mayer’s hematoxylin (Dako, #S330930-2, Germany) followed by bluing buffer (Dako, #CS70230-2, Germany). Sections were then counterstained with eosin (Sigma-Aldrich, #HT110216, Germany), mounted in 85% glycerol (Merck Millipore, #8187091000, Germany), and covered with a glass coverslip.

To visualize collagen accumulation during chronic liver disease progression 7-d after infection, Sirius Red (SR) staining was performed using a commercially available kit (Polyscience Europe GmbH, Germany). Slides were digitized at 40× magnification using a Hamamatsu L11600 slide scanner equipped with NDP.view2 Plus software (version U12388 02). Whole slide images were imported into QuPath (v0.6) (*64*). Liver parenchyma was annotated using the tissue detection function, with parameters adjusted to exclude image borders and large vessels. Collagen positive and total tissue areas were quantified using optimized staining vectors for the red collagen signal, a pixel resolution of 1.82 µm/pixel, Gaussian smoothing (sigma = 2), and a threshold of 0.14. The relative collagen area fraction (collagen positive area/total tissue area) was exported and compared.

### Regression model setup

The predictive performance of circulating IL-6 levels for 7-d outcomes was evaluated at serial time points (6, 24, and 48 h after infection) using receiver operating characteristic (ROC) curve analysis and calculation of area under the curve (AUC). Outcome prediction was modeled by logistic regression using pROC (R v1.19.0.1) (*65*). Based on model performance at 24 h after infection, an optimal circulating IL-6 cutoff of 15,052 pg µL^-1^ was defined to stratify septic mice into predicted survivors and predicted non-survivors (*44, 66*).

For subsequent analyses, mice from all experimental conditions with circulating IL-6 levels above this threshold at 24 h were classified as predicted non-survivors. Application of this classification to the 24 h cohort yielded the following distribution of predicted death: non-septic mice, 0/24; low-pathogen load group: vehicle-treated septic mice, 7/12, and Danirixin-treated septic mice, 5/12; high-pathogen load group: vehicle-treated septic mice, 11/12 and Danirixin-treated septic mice, 10/12.

To develop a clinically accessible classifier for the defined transcriptomic MTS subtypes, prediction of transcriptomic MTS from 24 h plasma IL 6 levels and initial pathogen load was performed using multinomial logistic regression. The full dataset included all 38 mice with available MTS labels and both biomarker measurements. The model was fitted in nnet (R v7.3-20), with MTS specified as the nominal outcome and IL 6 and PCI volume included as continuous linear predictors without interaction terms. For each animal, the predicted probability of MTS was calculated.

### Statistical analyses

All statistical analyses were performed in R (v4.5.1). Sample sizes were determined by power analyses based on prior experimental estimates. The primary hypothesis was to assess the efficacy of Danirixin relative to vehicle control. Experiments were conducted across multiple independent runs, with interim ethical evaluations after each iteration to ensure responsible animal use. For survival studies, sample size was calculated to distinguish moderate sepsis (48 h mortality: 20%) from severe sepsis (48 h mortality: 50%). To detect an absolute mortality difference of 30% with a power ≥ 0.8 at α = 0.05, a two-sided log-rank test was assumed, with a hazard ratio of 2.5, equal accrual and follow-up times (2 units each), and no subject attrition.

Under these assumptions, 47 animals per group were required. For analyses of immune response at 24 h after infection, MCP-1 was defined as the primary endpoint. Power analysis for a minimum 10% intergroup difference under nonparametric assumptions indicated a required sample size of 12 mice per group. Simulations based on Kruskal-Wallis test confirmed an all-pairs power of 0.8 and an any-pairs power of 1.

Data normality was assessed using the Shapiro-Wilk test. Non-normally distributed data were presented as median ± interquartile range (IQR) and were compared using the Wilcoxon rank-sum test or Kruskal-Wallis test, as appropriate. P-values from multiple comparisons were adjusted using the Holm-Bonferroni method. Survival distributions were estimated by the Kaplan-Meier method and compared using the log-rank test in survival (R v3.8-3) (*67*) and survminer (R v0.5.0). Correlations were assessed using Spearman rank correlation with 95% confidence intervals and visualized in corrplot (R v0.95).

PCA was applied to multiple independent datasets, including longitudinal systemic inflammatory profiles at 7-d after infection (CXCL1, CXCL2, IL-6), compartment-specific flow cytometric immune infiltration and inflammatory parameters at 24 h after infection (IL-6, CFUs, and NETosis), as well as matched bulk RNA-seq data from the blood and liver. Continuous variables were mean-centered and scaled to unit variance by Z-score normalization and PCA was computed using singular value decomposition. For statistical assessment of clustering in PCA space, individual sample scores for the first and second principal components (PC1 and PC2) were extracted.

Two-group comparisons for therapeutic intervention (vehicle vs. Danirixin), infectious gradients (sham-, low- and high-pathogen load, as applicable), and outcome (survivors vs. non-survivors or predicted survivors vs. predicted non-survivors) were performed using the Wilcoxon rank-sum test. Multiple-group comparisons for the human sepsis cohort (healthy, sepsis CTS patients) were performed using the Wilcoxon test followed by Holm-Bonferroni correction. Data visualization was performed in ggplot2 (R v4.0.2) and ggpubr (R v0.6.0). A two-sided p-value < 0.05 was considered statistically significant (ns, not significant; * p < 0.05; ** p < 0.01; *** p < 0.001; **** p < 0.0001).

## List of Supplementary Materials

Fig. S1 to S6

## Acknowledgments

The authors thank Dr. Jessica Barth, Dr. Ling Xiong, Francesca Kasimir, Masoumeh Eshaghi, and Christine Weiler from Jena University Hospital for technical assistance with animal experiments, bacterial quantification and histological section preparation for imaging. We also gratefully acknowledge Dr. Andrew J. Kwok, Prof. Julian C. Knight, their teams, and the Emergency Medicine Research Oxford (EMROx) consortium for making their omics data publicly available for reuse. During manuscript preparation, the authors used ChatGPT-5.4 for language polishing and grammar review. The authors take full responsibility for the content of the manuscript.

## Author contributions

Conceptualization: U.D., N.G., B.P.S., M.B., A.T.P.

Methodology: N.L., J.H., M.A., A.T.P.

Investigation: N.L., J.H., M.A., N.G., A.T.P.

Visualization: N.L., J.H., M.A., A.T.P.

Funding acquisition: M.B., A.T.P.

Project administration: M.B., A.T.P.

Supervision: B.P.S., M.B., A.T.P.

Writing – original draft: N.L., A.T.P.

Writing – review & editing: N.L., J.H., M.A., U.D., N.G., B.P.S., M.B., A.T.P.

## Data and materials availability

The original mRNA sequencing datasets generated and analyzed in this study have been deposited in the Zenodo repository under DOI: 10.5281/zenodo.19496937. Publicly available human datasets re-analyzed in this study were also obtained from Zenodo for the CITE-seq dataset (DOI: 10.5281/zenodo.7723202), deposited by the Emergency Medicine Research Oxford (EMROx) consortium.

## References

1. M. Shankar-Hari, T. Calandra, M. P. Soares, M. Bauer, W. J. Wiersinga, H. C. Prescott, J. C. Knight, K. J. Baillie, L. D. J. Bos, L. P. G. Derde, S. Finfer, R. S. Hotchkiss, J. Marshall, P. J. M. Openshaw, C. W. Seymour, F. Venet, J. L. Vincent, C. Le Tourneau, A. H. Maitland-van der Zee, I. B. McInnes, T. van der Poll, Reframing sepsis immunobiology for translation: towards informative subtyping and targeted immunomodulatory therapies. Lancet Respir Med, (2024).

2. B. P. Scicluna, K. Cano-Gamez, K. L. Burnham, E. E. Davenport, A. R. Moore, S. Khan, C. J. Hinds, O. L. Cremer, P. Khatri, T. E. Sweeney, J. C. Knight, T. van der Poll, A consensus blood transcriptomic framework for sepsis. Nat Med 31, 4119–4130 (2025).

3. A. J. Kwok, A. Allcock, R. C. Ferreira, E. Cano-Gamez, M. Smee, K. L. Burnham, Y. X. Zurke, S. McKechnie, A. J. Mentzer, C. Monaco, I. A. Udalova, C. J. Hinds, J. A. Todd, E. E. Davenport, J. C. Knight, Neutrophils and emergency granulopoiesis drive immune suppression and an extreme response endotype during sepsis. Nat Immunol 24, 767–779 (2023).

4. T. J. LaSalle, A. L. K. Gonye, S. S. Freeman, P. Kaplonek, I. Gushterova, K. R. Kays, K. Manakongtreecheep, J. Tantivit, M. Rojas-Lopez, B. C. Russo, N. Sharma, M. F. Thomas, K. M. Lavin-Parsons, B. M. Lilly, B. N. McKaig, N. C. Charland, H. K. Khanna, C. L. Lodenstein, J. D. Margolin, E. M. Blaum, P. B. Lirofonis, O. Y. Revach, A. Mehta, A. Sonny, R. P. Bhattacharyya, B. A. Parry, M. B. Goldberg, G. Alter, M. R. Filbin, A. C. Villani, N. Hacohen, M. Sade-Feldman, Longitudinal characterization of circulating neutrophils uncovers phenotypes associated with severity in hospitalized COVID-19 patients. Cell Rep Med 3, 100779 (2022).

5. W. Bao, H. Xing, S. Cao, X. Long, H. Liu, J. Ma, F. Guo, Z. Deng, X. Liu, Neutrophils restrain sepsis associated coagulopathy via extracellular vesicles carrying superoxide dismutase 2 in a murine model of lipopolysaccharide induced sepsis. Nat Commun 13, 4583 (2022).

6. X. Qi, Y. Yu, R. Sun, J. Huang, L. Liu, Y. Yang, T. Rui, B. Sun, Identification and characterization of neutrophil heterogeneity in sepsis. Crit Care 25, 50 (2021).

7. N. Liu, M. Bauer, A. T. Press, The immunological function of CXCR2 in the liver during sepsis. J Inflamm (Lond) 19, 23 (2022).

8. K. J. Eash, A. M. Greenbaum, P. K. Gopalan, D. C. Link, CXCR2 and CXCR4 antagonistically regulate neutrophil trafficking from murine bone marrow. J Clin Invest 120, 2423–2431 (2010).

9. P. Delobel, B. Ginter, E. Rubio, K. Balabanian, G. Lazennec, CXCR2 intrinsically drives the maturation and function of neutrophils in mice. Front Immunol 13, 1005551 (2022).

10. J. Zhang, Y. Shao, J. Wu, J. Zhang, X. Xiong, J. Mao, Y. Wei, C. Miao, H. Zhang, Dysregulation of neutrophil in sepsis: recent insights and advances. Cell Commun Signal 23, 87 (2025).

11. Polyploidy in liver development, homeostasis and disease.

12. A. Charoensappakit, K. Sae-Khow, N. Vutthikraivit, P. Maneesow, T. Sriprasart, M. Pachinburavan, A. Leelahavanichkul, Immune suppressive activities of low-density neutrophils in sepsis and potential use as a novel biomarker of sepsis-induced immune suppression. Sci Rep 15, 9458 (2025).

13. C. Seree-Aphinan, P. Vichitkunakorn, R. Navakanitworakul, B. Khwannimit, Distinguishing Sepsis From Infection by Neutrophil Dysfunction: A Promising Role of CXCR2 Surface Level. Front Immunol 11, 608696 (2020).

14. M. Alsabani, S. T. Abrams, Z. Cheng, B. Morton, S. Lane, S. Alosaimi, W. Yu, G. Wang, C. H. Toh, Reduction of NETosis by targeting CXCR1/2 reduces thrombosis, lung injury, and mortality in experimental human and murine sepsis. Br J Anaesth 128, 283–293 (2022).

15. T. L. Ness, C. M. Hogaboam, R. M. Strieter, S. L. Kunkel, Immunomodulatory role of CXCR2 during experimental septic peritonitis. J Immunol 171, 3775–3784 (2003).

16. T. Jamieson, M. Clarke, C. W. Steele, M. S. Samuel, J. Neumann, A. Jung, D. Huels, M. F. Olson, S. Das, R. J. Nibbs, O. J. Sansom, Inhibition of CXCR2 profoundly suppresses inflammation-driven and spontaneous tumorigenesis. J Clin Invest 122, 3127–3144 (2012).

17. C. Montemagno, A. Jacquel, C. Pandiani, O. Rastoin, R. Dawaliby, T. Schmitt, M. Bourgoin, H. Palenzuela, A. L. Rossi, D. Ambrosetti, J. Durivault, F. Luciano, D. Borchiellini, J. Le Du, L. C. P. Gonçalves, P. Auberger, R. Benhida, L. Kinget, B. Beuselinck, C. Ronco, G. Pagès, M. Dufies, Unveiling CXCR2 as a promising therapeutic target in renal cell carcinoma: exploring the immunotherapeutic paradigm shift through its inhibition by RCT001. J Exp Clin Cancer Res 43, 86 (2024).

18. H. R. Keir, H. Richardson, C. Fillmore, A. Shoemark, A. L. Lazaar, B. E. Miller, R. Tal-Singer, J. D. Chalmers, D. Mohan, CXCL-8-dependent and -independent neutrophil activation in COPD: experiences from a pilot study of the CXCR2 antagonist danirixin. ERJ Open Res 6, (2020).

19. A. L. Lazaar, B. E. Miller, A. C. Donald, T. Keeley, C. Ambery, J. Russell, H. Watz, R. Tal-Singer, CXCR2 antagonist for patients with chronic obstructive pulmonary disease with chronic mucus hypersecretion: a phase 2b trial. Respir Res 21, 149 (2020).

20. E. E. Davenport, K. L. Burnham, J. Radhakrishnan, P. Humburg, P. Hutton, T. C. Mills, A. Rautanen, A. C. Gordon, C. Garrard, A. V. Hill, C. J. Hinds, J. C. Knight, Genomic landscape of the individual host response and outcomes in sepsis: a prospective cohort study. Lancet Respir Med 4, 259–271 (2016).

21. A. R. Moore, H. Zheng, A. Ganesan, Y. Hasin-Brumshtein, M. V. Maddali, J. E. Levitt, T. van der Poll, J. Lu, H. R. Bouma, B. P. Scicluna, E. J. Giamarellos-Bourboulis, A. Kotsaki, I. Martin-Loeches, A. Garduno, J. Hinson, R. E. Rothman, J. Sevransky, D. W. Wright, M. R. Atreya, L. L. Moldawer, P. A. Efron, M. Kralovcova, T. Karvunidis, H. M. Giannini, N. J. Meyer, T. E. Sweeney, A. J. Rogers, P. Khatri, A consensus immune dysregulation framework for sepsis and critical illnesses. Nat Med 31, 4084–4096 (2025).

22. I. Rumienczyk, M. Kulecka, J. Ostrowski, D. Mar, K. Bomsztyk, S. W. Standage, M. Mikula, Multi-Organ Transcriptome Dynamics in a Mouse Model of Cecal Ligation and Puncture-Induced Polymicrobial Sepsis. J Inflamm Res 14, 2377–2388 (2021).

23. G. Lu, F. Han, Y. Wang, C. Yuan, Q. Zhu, T. Xia, L. Chen, X. Dong, Y. Ding, W. Xiao, Y. Zhang, J. Pan, H. Xu, W. Chen, B. Tu, W. Li, F. Wang, W. Gong, L. Hu, Src Reduces Neutrophil Extracellular Traps Generation and Resolves Acute Organ Damage. Adv Sci (Weinh) 12, e06028 (2025).

24. R. Zhang, L. Chen, W. Xu, J. Ye, Z. Wang, W. Xu, B. Wu, CXCR2(+) Neutrophils Drive Neutrophil Extracellular Traps Formation and Exacerbate Pulpitis in Rats: An In Vitro and In Vivo Laboratory Investigation. *Int Endod J*, (2026).

25. K. Liu, L. Shen, M. Wu, Z. J. Liu, T. Hua, Structural insights into the activation of chemokine receptor CXCR2. Febs j 289, 386–393 (2022).

26. K. Liu, L. Wu, S. Yuan, M. Wu, Y. Xu, Q. Sun, S. Li, S. Zhao, T. Hua, Z. J. Liu, Structural basis of CXC chemokine receptor 2 activation and signalling. Nature 585, 135–140 (2020).

27. F. Rios-Santos, J. C. Alves-Filho, F. O. Souto, F. Spiller, A. Freitas, C. M. Lotufo, M. B. Soares, R. R. Dos Santos, M. M. Teixeira, F. Q. Cunha, Down-regulation of CXCR2 on neutrophils in severe sepsis is mediated by inducible nitric oxide synthase-derived nitric oxide. Am J Respir Crit Care Med 175, 490–497 (2007).

28. R. Sun, J. Huang, H. Jin, X. Wen, X. Gao, B. Sun, NETosis-Dependent Generation of Immunodeficient Low-Density Neutrophils Exacerbates Sepsis-Induced Acute Lung Injury. Int J Mol Sci 27, (2026).

29. Á. Teijeira, S. Garasa, M. Gato, C. Alfaro, I. Migueliz, A. Cirella, C. de Andrea, M. C. Ochoa, I. Otano, I. Etxeberria, M. P. Andueza, C. P. Nieto, L. Resano, A. Azpilikueta, M. Allegretti, M. de Pizzol, M. Ponz-Sarvisé, A. Rouzaut, M. F. Sanmamed, K. Schalper, M. Carleton, M. Mellado, M. E. Rodriguez-Ruiz, P. Berraondo, J. L. Perez-Gracia, I. Melero, CXCR1 and CXCR2 Chemokine Receptor Agonists Produced by Tumors Induce Neutrophil Extracellular Traps that Interfere with Immune Cytotoxicity. Immunity 52, 856–871.e858 (2020).

30. W. Jing, G. Wang, Z. Cui, X. Li, S. Zeng, X. Jiang, W. Li, B. Han, N. Xing, Y. Zhao, S. Chen, B. Shi, Tumor-neutrophil cross talk orchestrates the tumor microenvironment to determine the bladder cancer progression. Proc Natl Acad Sci U S A 121, e2312855121 (2024).

31. N. C. Kaneider, A. Agarwal, A. J. Leger, A. Kuliopulos, Reversing systemic inflammatory response syndrome with chemokine receptor pepducins. Nat Med 11, 661–665 (2005).

32. V. Wieser, T. E. Adolph, B. Enrich, A. Kuliopulos, A. Kaser, H. Tilg, N. C. Kaneider, Reversal of murine alcoholic steatohepatitis by pepducin-based functional blockade of interleukin-8 receptors. Gut 66, 930–938 (2017).

33. H. L. Van Sweringen, N. Sakai, R. C. Quillin, J. Bailey, R. Schuster, J. Blanchard, H. Goetzman, C. C. Caldwell, M. J. Edwards, A. B. Lentsch, Roles of hepatocyte and myeloid CXC chemokine receptor-2 in liver recovery and regeneration after ischemia/reperfusion in mice. Hepatology 57, 331–338 (2013).

34. P. Opfermann, U. Derhaschnig, A. Felli, J. Wenisch, D. Santer, A. Zuckermann, M. Dworschak, B. Jilma, B. Steinlechner, A pilot study on reparixin, a CXCR1/2 antagonist, to assess safety and efficacy in attenuating ischaemia-reperfusion injury and inflammation after on-pump coronary artery bypass graft surgery. Clin Exp Immunol 180, 131–142 (2015).

35. N. H. Mohamad Zaki, J. Shiota, A. N. Calder, T. M. Keeley, B. L. Allen, K. Nakao, L. C. Samuelson, N. Razumilava, C-X-C motif chemokine ligand 1 induced by Hedgehog signaling promotes mouse extrahepatic bile duct repair after acute injury. Hepatology 76, 936–950 (2022).

36. D. Ye, K. Yang, S. Zang, Z. Lin, H. T. Chau, Y. Wang, J. Zhang, J. Shi, A. Xu, S. Lin, Y. Wang, Lipocalin-2 mediates non-alcoholic steatohepatitis by promoting neutrophil-macrophage crosstalk via the induction of CXCR2. J Hepatol 65, 988–997 (2016).

37. T. Konishi, R. M. Schuster, H. S. Goetzman, C. C. Caldwell, A. B. Lentsch, Cell-specific regulatory effects of CXCR2 on cholestatic liver injury. Am J Physiol Gastrointest Liver Physiol 317, G773–g783 (2019).

38. Y. Hao, T. Stuart, M. H. Kowalski, S. Choudhary, P. Hoffman, A. Hartman, A. Srivastava, G. Molla, S. Madad, C. Fernandez-Granda, R. Satija, Dictionary learning for integrative, multimodal and scalable single-cell analysis. Nat Biotechnol 42, 293–304 (2024).

39. Y. Hua, L. Weng, F. Zhao, F. Rambow, SeuratExtend: streamlining single-cell RNA-seq analysis through an integrated and intuitive framework. Gigascience 14, (2025).

40. N. Liu, M. Sonawane, O. Sommerfeld, C. M. Svensson, M. T. Figge, R. Bauer, S. J. Bischoff, M. Bauer, M. F. Osuchowski, A. T. Press, Metamizole outperforms meloxicam in sepsis: insights on analgesics, survival and immunomodulation in the peritoneal contamination and infection sepsis model. Front Immunol 15, 1432307 (2024).

41. L. Xiong, D. Beyer, N. Liu, T. Lehmann, S. Neugebauer, S. Schaeuble, O. Sommerfeld, P. Ernst, C. M. Svensson, S. Nietzsche, S. Scholl, T. Bruns, N. Gaßler, M. H. Gräler, M. T. Figge, G. Panagiotou, M. Bauer, A. T. Press, Targeting protein kinase C-α prolongs survival and restores liver function in sepsis: Evidence from preclinical models. Pharmacol Res 212, 107581 (2025).

42. M. F. Osuchowski, A. Ayala, S. Bahrami, M. Bauer, M. Boros, J. M. Cavaillon, I. H. Chaudry, C. M. Coopersmith, C. S. Deutschman, S. Drechsler, P. Efron, C. Frostell, G. Fritsch, W. Gozdzik, J. Hellman, M. Huber-Lang, S. Inoue, S. Knapp, A. V. Kozlov, C. Libert, J. C. Marshall, L. L. Moldawer, P. Radermacher, H. Redl, D. G. Remick, M. Singer, C. Thiemermann, P. Wang, W. J. Wiersinga, X. Xiao, B. Zingarelli, Minimum Quality Threshold in Pre-Clinical Sepsis Studies (MQTiPSS): An International Expert Consensus Initiative for Improvement of Animal Modeling in Sepsis. Shock 50, 377–380 (2018).

43. N. Percie du Sert, V. Hurst, A. Ahluwalia, S. Alam, M. T. Avey, M. Baker, W. J. Browne, A. Clark, I. C. Cuthill, U. Dirnagl, M. Emerson, P. Garner, S. T. Holgate, D. W. Howells, N. A. Karp, S. E. Lazic, K. Lidster, C. J. MacCallum, M. Macleod, E. J. Pearl, O. H. Petersen, F. Rawle, P. Reynolds, K. Rooney, E. S. Sena, S. D. Silberberg, T. Steckler, H. Würbel, The ARRIVE guidelines 2.0: Updated guidelines for reporting animal research. PLoS Biol 18, e3000410 (2020).

44. M. F. Osuchowski, K. Welch, J. Siddiqui, D. G. Remick, Circulating cytokine/inhibitor profiles reshape the understanding of the SIRS/CARS continuum in sepsis and predict mortality. J Immunol 177, 1967–1974 (2006).

45. M. F. Osuchowski, J. Connett, K. Welch, J. Granger, D. G. Remick, Stratification is the key: inflammatory biomarkers accurately direct immunomodulatory therapy in experimental sepsis. Crit Care Med 37, 1567–1573 (2009).

46. P. Raeven, S. Drechsler, K. M. Weixelbaumer, D. Bastelica, F. Peiretti, A. Klotz, M. Jafarmadar, H. Redl, S. Bahrami, M. C. Alessi, P. J. Declerck, M. F. Osuchowski, Systemic inhibition and liver-specific over-expression of PAI-1 failed to improve survival in all-inclusive populations or homogenous cohorts of CLP mice. J Thromb Haemost 12, 958–969 (2014).

47. C. W. Meyer, Y. Ootsuka, A. A. Romanovsky, Body Temperature Measurements for Metabolic Phenotyping in Mice. Front Physiol 8, 520 (2017).

48. H. Xiao, J. Siddiqui, D. G. Remick, Mechanisms of Mortality in Early and Late Sepsis. Infection and Immunity 74, 5227–5235 (2006).

49. M. F. Osuchowski, K. Welch, J. Siddiqui, D. G. Remick, Circulating Cytokine/Inhibitor Profiles Reshape the Understanding of the SIRS/CARS Continuum in Sepsis and Predict Mortality1. The Journal of Immunology 177, 1967–1974 (2006).

50. E. Li, X. Yang, Y. Du, G. Wang, D. W. Chan, D. Wu, P. Xu, P. Ni, D. Xu, Y. Hu, CXCL8 Associated Dendritic Cell Activation Marker Expression and Recruitment as Indicators of Favorable Outcomes in Colorectal Cancer. Front Immunol 12, 667177 (2021).

51. A. M. Bolger, M. Lohse, B. Usadel, Trimmomatic: a flexible trimmer for Illumina sequence data. Bioinformatics 30, 2114–2120 (2014).

52. A. Dobin, C. A. Davis, F. Schlesinger, J. Drenkow, C. Zaleski, S. Jha, P. Batut, M. Chaisson, T. R. Gingeras, STAR: ultrafast universal RNA-seq aligner. Bioinformatics 29, 15–21 (2013).

53. Y. Liao, G. K. Smyth, W. Shi, featureCounts: an efficient general purpose program for assigning sequence reads to genomic features. Bioinformatics 30, 923–930 (2013).

54. S. C. Dyer, O. Austine-Orimoloye, A. G. Azov, M. Barba, I. Barnes, V. P. Barrera-Enriquez, A. Becker, R. Bennett, M. Beracochea, A. Berry, J. Bhai, S. K. Bhurji, S. Boddu, P. R. Branco Lins, L. Brooks, S. B. Ramaraju, L. I. Campbell, M. C. Martinez, M. Charkhchi, L. A. Cortes, C. Davidson, S. Denni, K. Dodiya, S. Donaldson, B. El Houdaigui, T. El Naboulsi, O. Falola, R. Fatima, T. Genez, J. G. Martinez, T. Gurbich, M. Hardy, Z. Hollis, T. Hunt, M. Kay, V. Kaykala, D. Lemos, D. Lodha, N. Mathlouthi, G. A. Merino, R. Merritt, L. P. Mirabueno, A. Mushtaq, S. N. Hossain, J. G. Pérez-Silva, M. Perry, I. Piližota, D. Poppleton, I. Prosovetskaia, S. Raj, A. Imran A. Salam, S. Saraf, N. Saraiva-Agostinho, S. Sinha, B. Sipos, V. Sitnik, E. Steed, M.-M. Suner, L. Surapaneni, K. Sutinen, F. F. Tricomi, I. Tsang, D. Urbina-Gómez, A. Veidenberg, T. A. Walsh, N. L. Willhoft, J. Allen, J. Alvarez-Jarreta, M. Chakiachvili, J. Cheema, J. B. da Rocha, N. H. De Silva, S. Giorgetti, L. Haggerty, G. R. Ilsley, J. Keatley, J. E. Loveland, B. Moore, J. M. Mudge, G. Naamati, J. Tate, S. J. Trevanion, A. Winterbottom, B. Flint, A. Frankish, S. E. Hunt, R. D. Finn, M. A. Freeberg, P. W. Harrison, F. J. Martin, A. D. Yates, Ensembl 2025. Nucleic Acids Research 53, D948–D957 (2024).

55. Y. Zhang, G. Parmigiani, W. E. Johnson, ComBat-seq: batch effect adjustment for RNA-seq count data. NAR Genomics and Bioinformatics 2, (2020).

56. M. I. Love, W. Huber, S. Anders, Moderated estimation of fold change and dispersion for RNA-seq data with DESeq2. Genome Biol 15, 550 (2014).

57. B. M. Schilder, A. E. Murphy, N. G. Skene, orthogene: a Bioconductor package to easily map genes within and across hundreds of species. *bioRxiv*, 2026.2001.2017.700094 (2026).

58. Z. Gu, Complex heatmap visualization. Imeta 1, e43 (2022).

59. A. A. Sergushichev, An algorithm for fast preranked gene set enrichment analysis using cumulative statistic calculation. bioRxiv, 060012 (2016).

60. A. Liberzon, A. Subramanian, R. Pinchback, H. Thorvaldsdóttir, P. Tamayo, J. P. Mesirov, Molecular signatures database (MSigDB) 3.0. Bioinformatics 27, 1739–1740 (2011).

61. G. Yu, L.-G. Wang, Y. Han, Q.-Y. He, clusterProfiler: an R Package for Comparing Biological Themes Among Gene Clusters. OMICS: A Journal of Integrative Biology 16, 284–287 (2012).

62. A. Cossarizza, H. D. Chang, A. Radbruch, S. Abrignani, R. Addo, M. Akdis, I. Andrä, F. Andreata, F. Annunziato, E. Arranz, P. Bacher, S. Bari, V. Barnaba, J. Barros-Martins, D. Baumjohann, C. G. Beccaria, D. Bernardo, D. A. Boardman, J. Borger, C. Böttcher, L. Brockmann, M. Burns, D. H. Busch, G. Cameron, I. Cammarata, A. Cassotta, Y. Chang, F. G. Chirdo, E. Christakou, L. Čičin-Šain, L. Cook, A. J. Corbett, R. Cornelis, L. Cosmi, M. S. Davey, S. De Biasi, G. De Simone, G. Del Zotto, M. Delacher, F. Di Rosa, J. Di Santo, A. Diefenbach, J. Dong, T. Dörner, R. J. Dress, C. A. Dutertre, S. B. G. Eckle, P. Eede, M. Evrard, C. S. Falk, M. Feuerer, S. Fillatreau, A. Fiz-Lopez, M. Follo, G. A. Foulds, J. Fröbel, N. Gagliani, G. Galletti, A. Gangaev, N. Garbi, J. A. Garrote, J. Geginat, N. A. Gherardin, L. Gibellini, F. Ginhoux, D. I. Godfrey, P. Gruarin, C. Haftmann, L. Hansmann, C. M. Harpur, A. C. Hayday, G. Heine, D. C. Hernández, M. Herrmann, O. Hoelsken, Q. Huang, S. Huber, J. E. Huber, J. Huehn, M. Hundemer, W. Y. K. Hwang, M. Iannacone, S. M. Ivison, H. M. Jäck, P. K. Jani, B. Keller, N. Kessler, S. Ketelaars, L. Knop, J. Knopf, H. F. Koay, K. Kobow, K. Kriegsmann, H. Kristyanto, A. Krueger, J. F. Kuehne, H. Kunze-Schumacher, P. Kvistborg, I. Kwok, D. Latorre, D. Lenz, M. K. Levings, A. C. Lino, F. Liotta, H. M. Long, E. Lugli, K. N. MacDonald, L. Maggi, M. K. Maini, F. Mair, C. Manta, R. A. Manz, M. F. Mashreghi, A. Mazzoni, J. McCluskey, H. E. Mei, F. Melchers, S. Melzer, D. Mielenz, L. Monin, L. Moretta, G. Multhoff, L. E. Muñoz, M. Muñoz-Ruiz, F. Muscate, A. Natalini, K. Neumann, L. G. Ng, A. Niedobitek, J. Niemz, L. N. Almeida, S. Notarbartolo, L. Ostendorf, L. J. Pallett, A. A. Patel, G. I. Percin, G. Peruzzi, M. Pinti, A. G. Pockley, K. Pracht, I. Prinz, I. Pujol-Autonell, N. Pulvirenti, L. Quatrini, K. M. Quinn, H. Radbruch, H. Rhys, M. B. Rodrigo, C. Romagnani, C. Saggau, S. Sakaguchi, F. Sallusto, L. Sanderink, I. Sandrock, C. Schauer, A. Scheffold, H. U. Scherer, M. Schiemann, F. A. Schildberg, K. Schober, J. Schoen, W. Schuh, T. Schüler, A. R. Schulz, S. Schulz, J. Schulze, S. Simonetti, J. Singh, K. M. Sitnik, R. Stark, S. Starossom, C. Stehle, F. Szelinski, L. Tan, A. Tarnok, J. Tornack, T. I. M. Tree, J. J. P. van Beek, W. van de Veen, K. van Gisbergen, C. Vasco, N. A. Verheyden, A. von Borstel, K. A. Ward-Hartstonge, K. Warnatz, C. Waskow, A. Wiedemann, A. Wilharm, J. Wing, O. Wirz, J. Wittner, J. H. M. Yang, J. Yang, Guidelines for the use of flow cytometry and cell sorting in immunological studies (third edition). Eur J Immunol 51, 2708–3145 (2021).

63. M. F. Osuchowski, J. Siddiqui, S. Copeland, D. G. Remick, Sequential ELISA to profile multiple cytokines from small volumes. J Immunol Methods 302, 172–181 (2005).

64. P. Bankhead, M. B. Loughrey, J. A. Fernandez, Y. Dombrowski, D. G. McArt, P. D. Dunne, S. McQuaid, R. T. Gray, L. J. Murray, H. G. Coleman, J. A. James, M. Salto-Tellez, P. W. Hamilton, QuPath: Open source software for digital pathology image analysis. Sci Rep 7, 16878 (2017).

65. X. Robin, N. Turck, A. Hainard, N. Tiberti, F. Lisacek, J. C. Sanchez, M. Müller, pROC: an open-source package for R and S+ to analyze and compare ROC curves. BMC Bioinformatics 12, 77 (2011).

66. T. Skirecki, S. Drechsler, A. Jeznach, G. Hoser, M. Jafarmadar, J. Kawiak, M. F. Osuchowski, An Early Myelosuppression in the Acute Mouse Sepsis Is Partly Outcome-Dependent. Front Immunol 12, 708670 (2021).

67. T. M. Therneau, “A Package for Survival Analysis in R,” (2026).

