## Supplemental Figures for "Sepsis Subtypes in Blood and Liver Define Precision Strategy for CXCR2 Blockade": Supplemental Figures.docx

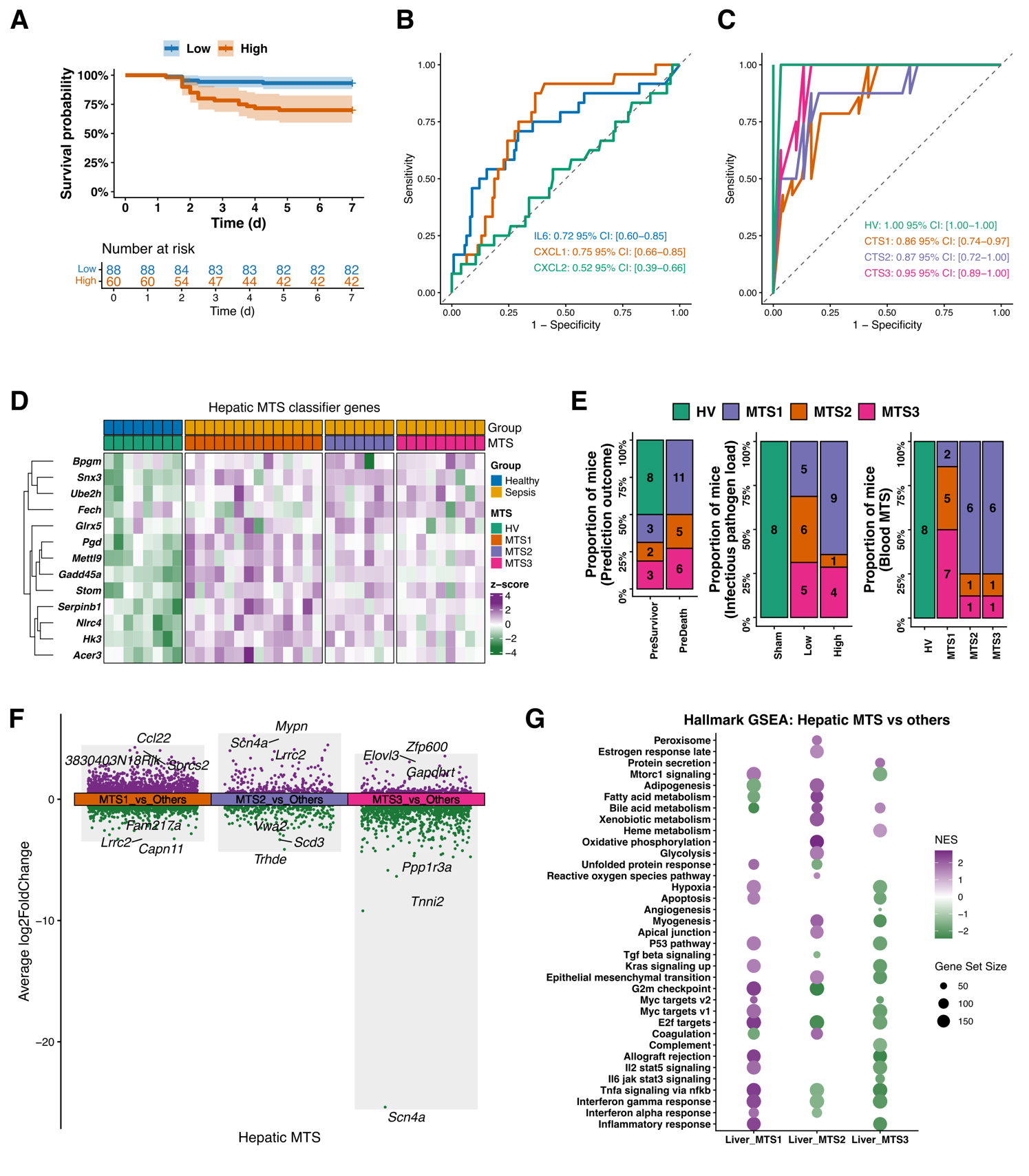

**Figure S1. Hepatic murine transcriptomic subtypes classification defines distinct clinical associations and functional programs in sepsis. (A)** Seven-day survival analysis of mice subjected to low- and high-pathogen-load sepsis with Kaplan–Meier curves. **(B)** ROC analysis evaluating the predictive performance of circulating CXCL1/2, and IL-6 levels for 7-d survival outcomes. **(C)** ROC analysis evaluating the accuracy of blood MTS model. **(D)** Heatmap of MTS classifier genes across 38 liver samples, including 8 healthy volunteers (HV), 14 MTS1, 7 MTS2, and 9 MTS3 samples. **(E)** Distribution of hepatic MTS subtypes stratified by predicted outcome, initial infectious pathogen load, and corresponding blood MTS classification. **(F)** DEGs analysis comparing each hepatic MTS subgroup against all remaining liver samples. **(G)** Hallmark GSEA performed for each hepatic MTS subtype using a one-vs-rest comparison strategy.

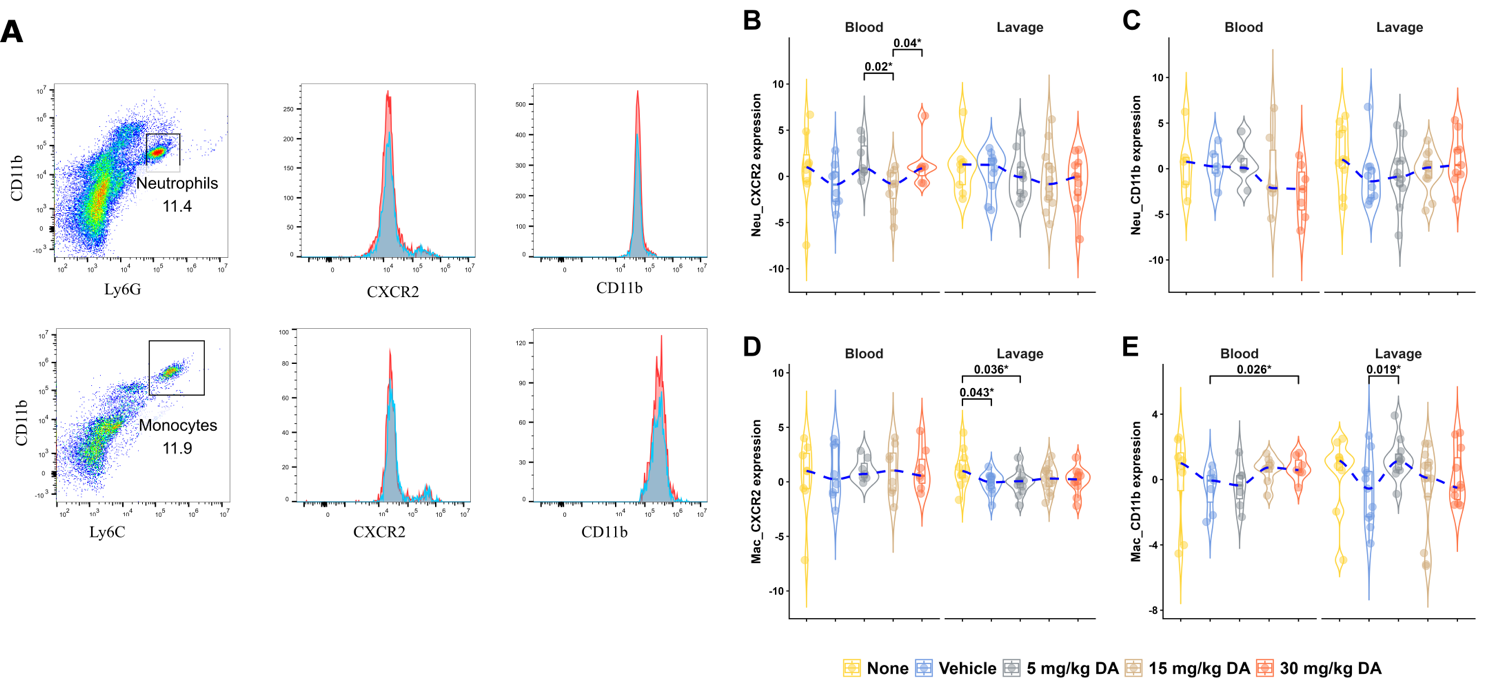

**Figure S2. Dose-dependent *in vivo* pharmacological profiling of CXCR2 antagonism by Danirixin. (A)** Representative flow cytometry gating strategy for identification of systemic and compartmental leukocyte populations. Leukocytes were identified as CD45⁺ live singlets, followed by specific delineation of neutrophils (CD11b⁺ Ly6G⁺), blood monocytes (CD11b⁺ Ly6C⁺), and peritoneal macrophages (CD11b⁺ F4/80⁺). Surface expression levels of CD11b and CXCR2 were quantified using gMFI. **(B-E)** Quantification of surface CXCR2 and CD11b expression (gMFI) on neutrophils and mononuclear phagocytes (macrophages/monocytes) isolated from whole blood and peritoneal lavage fluid (PLF). Data are presented as median ± IQR with asterisks denoting statistical significance as *p < 0.05.

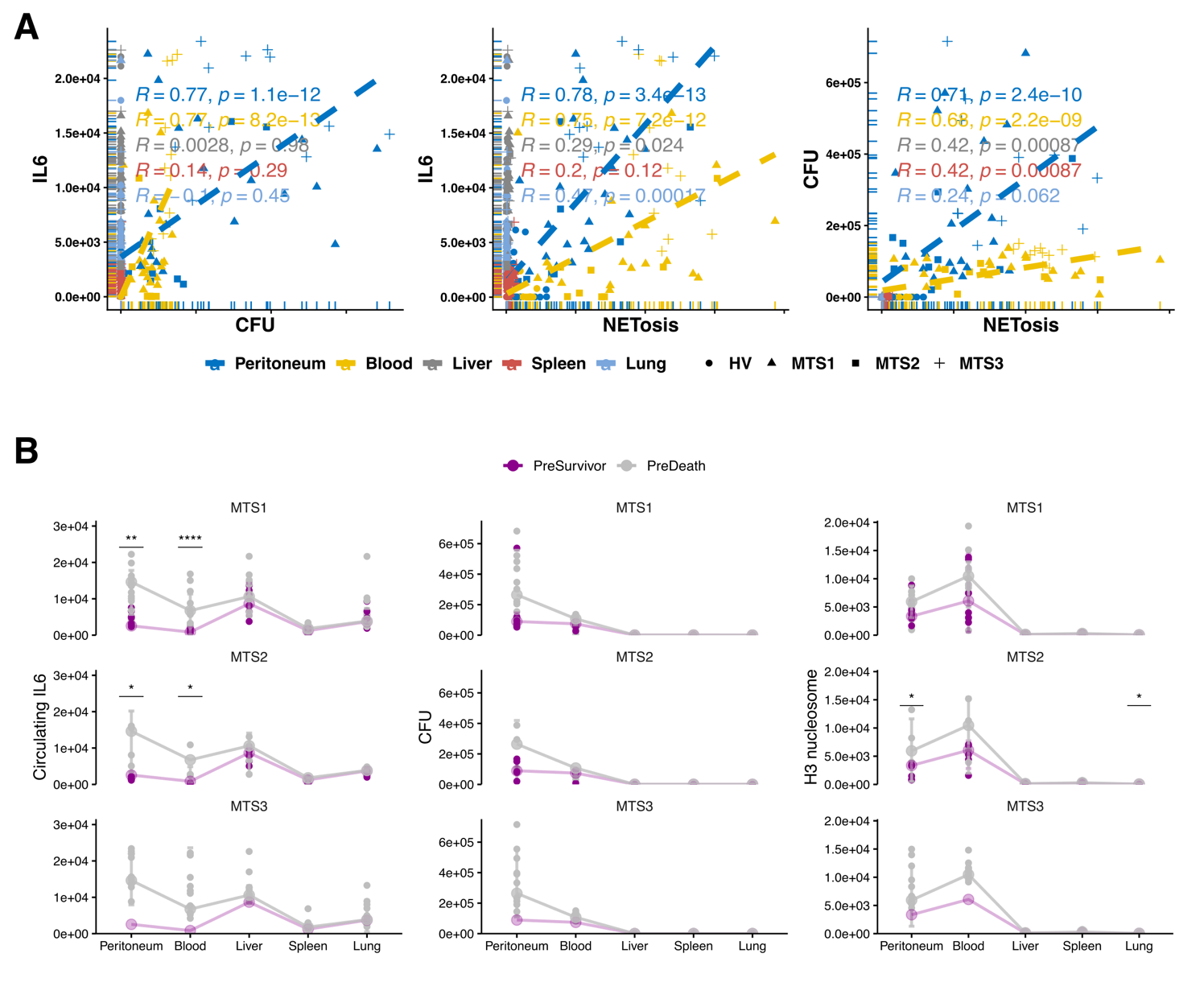

**Figure S3. Anatomical compartmentalization and trajectory mapping of inflammatory and pathological markers. (A)** Spearman correlation analyses of localized IL-6 levels, bacterial burden (CFU), and NETosis (Histone H3 nucleosomes) across the compartments 24 h after infection. **(B)** Cross-organ gradient mapping of IL-6, CFU, and H3 nucleosomes in MTS models. Asterisks denote statistical significance (ns, not significant; * p < 0.05; ** p < 0.01; *** p < 0.001; **** p < 0.0001).

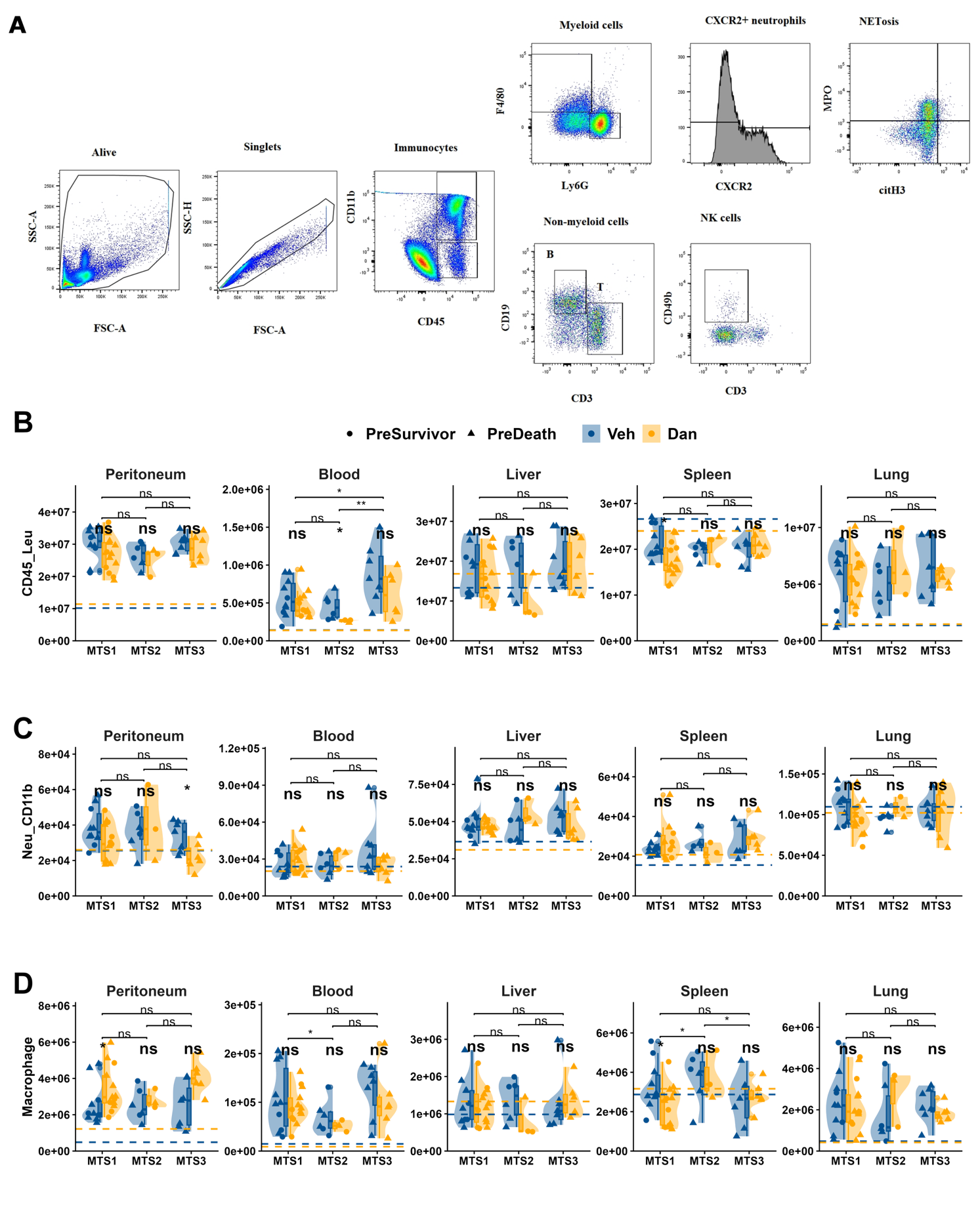

**Figure S4. Flow cytometric characterization and multi-organ immune profiling during polymicrobial sepsis. (A)** Representative gating strategy for identifying immune cell populations across tissues. Live CD45⁺ leukocytes are subdivided into myeloid (CD11b^+^ CD45⁺) and non-myeloid (CD11b⁻ CD45⁺) lineages. Myeloid cells are classified into tissue macrophages or blood monocytes (F4/80⁺ and/or Ly6C⁺) and neutrophils (Ly6G⁺), Neutrophils are subsequently evaluated for surface CXCR2 and CD11b expression. NETosis is identified by CitH3^+^ MPO⁺ cells. Non-myeloid cells are subdivided into T cells (CD3⁺ CD19⁻), B cells (CD19⁺ CD3⁻), and NK cells (CD49b⁺ CD3⁻). (**B-D**) Violin plot of (B) total CD45⁺ immune cell abundance, (C) neutrophil surface CD11b expression (geometrix mean fluorescence intensity, gMFI), and (D) blood monocyte and tissue macrophage accumulation, across tissues. Asterisks denote statistical significance ns, not significant; * p < 0.05; ** p < 0.01; *** p < 0.001; **** p < 0.0001).

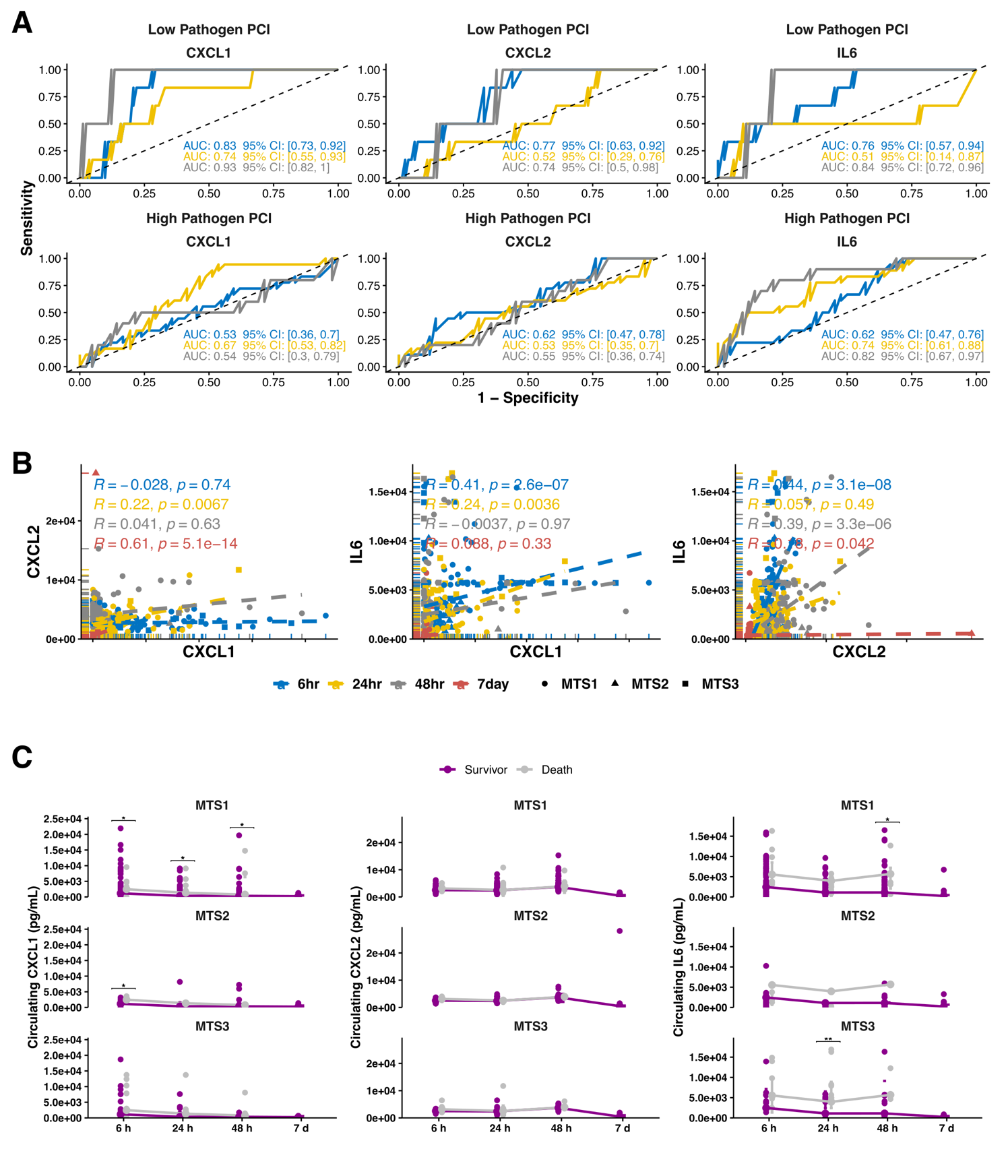

**Figure S5. Temporal dynamics of circulating CXCL1/2 and IL-6, correlation and outcome-associated predictive capacity in polymicrobial sepsis. (A)** ROC curves evaluating the capacity of circulating CXCL1/2, and IL‑6 to predict 7‑d mortality, stratified by pathogen load. AUC with 95% confidence intervals is indicated for each cytokine–time–pathogen combination. **(B)** Spearman cross‑correlations of circulating CXCL1/2, and IL‑6 concentrations across four time points after infection. **(C)** Longitudinal profiles of circulating CXCL1/2, and IL‑6 over 7 d, stratified by identified MTS and 7‑d survival outcome. Asterisks denote statistical significance (ns, not significant; * p < 0.05; ** p < 0.01; *** p < 0.001; **** p < 0.0001).

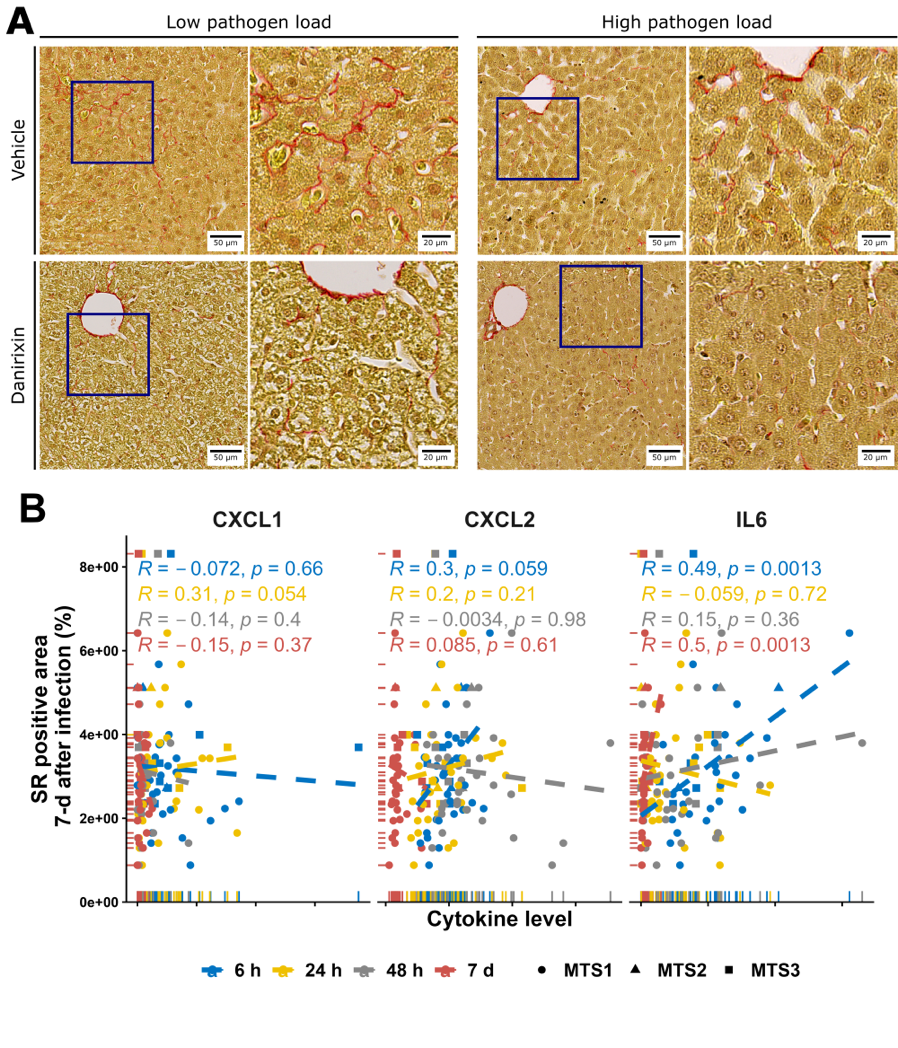

**Figure S6. Hepatic collagen deposition and its correlation with systemic CXCL1/2, and IL‑6 over time. (A)** Sirius Red (SR) staining of liver sections at 7 d after infection. Representative images and quantification of collagen deposition are shown, with samples stratified by initial pathogen load and treatment. **(B)** Sequential correlation analysis between hepatic collagen deposition and circulating concentrations of CXCL1/2, and IL‑6 measured at four time points after infection.
